# A small molecule that binds an RNA repeat expansion stimulates its decay via the exosome complex

**DOI:** 10.1101/2020.05.11.088427

**Authors:** Alicia J. Angelbello, Raphael I. Benhamou, Suzanne G. Rzuczek, Shruti Choudhary, Zhenzhi Tang, Jonathan L. Chen, Madhuparna Roy, Kye Won Wang, Ilyas Yildirim, Albert S. Jun, Charles A. Thornton, Matthew D. Disney

**Author notes:** These authors contributed equally to this work (Alicia J. Angelbello, Raphael I. Benhamou, Suzanne G. Rzuczek).

## Abstract

We describe the design of a small molecule that binds the structure of a r(CUG) repeat expansion [r(CUG)^exp^] and reverses molecular defects in two diseases mediated by the RNA - myotonic dystrophy type 1 (DM1) and Fuchs endothelial corneal dystrophy (FECD). Thus, a single structure-specific ligand has potential therapeutic benefit for multiple diseases, in contrast to oligonucleotide-based modalities that are customized for each disease by nature of targeting the gene that harbors the repeat. Indeed, the small molecule binds the target with nanomolar affinity and >100-fold specificity vs. many other RNAs and DNA. Interestingly, the compound’s downstream effects are different in the two diseases, owing to the location of the repeat expansion. In DM1, r(CUG)^exp^ is harbored in the 3’ untranslated region (UTR) of and mRNA, and the compound has no effect on the RNA’s abundance. In FECD, however, r(CUG)^exp^ is located in an intron, and the small molecule, by binding the repeat expansion, facilitates excision of the intron, which is then degraded by the exosome complex exonuclease, hRRP6. Thus, structure-specific, RNA-targeting small molecules can act disease-specifically to affect biology, either by disabling its gain-of-function mechanism (DM1) or by stimulating quality control pathways to rid a disease-affected cell of a toxic RNA (FECD).

**Significance statement:** Many different diseases are caused by toxic structured RNAs. Herein, we designed a lead small molecule that binds a toxic structure and rescues disease biology. We show that a structure-specific small molecule can improve disease-associated defects in two diseases that share the common toxic RNA structure. In one disease, the toxic structure is harbored in an intron and causes its retention. The compound facilitates processing of a retained intron, enabling the disease-affected cell to remove the toxic RNA.

## Introduction

Over 40 diseases are caused by RNA repeat expansions including Huntington’s disease (HD) (1), C9orf72-mediated amyotrophic lateral sclerosis/frontotemporal dementia (c9ALS/FTD) (2), myotonic dystrophy type 1 (DM1) (3), and Fuchs endothelial corneal dystrophy (FECD) (4). DM1 is caused by expression of an r(CUG) repeat expansion [r(CUG)^exp^] harbored in the 3’ untranslated region (UTR) of the *dystrophia myotonica protein kinase* (*DMPK*) mRNA (Fig. 1A) (3). DM1 is the most common form of adult-onset muscular dystrophy, affecting approximately 1 in 8,000 people, and r(CUG)^exp^ causes disease by a gain-of-function mechanism in which the repeat expansions binds to and sequesters various proteins, particularly the pre-mRNA splicing regulator muscleblind-like 1 (MBNL1), in nuclear foci (5). Sequestration of MBL1 by r(CUG)^exp^ limits the amount of MBNL1 available to regulate pre-mRNA splicing, causing system-wide defects that manifest themselves in DM1 (Fig. 1B) (6,7).

**Fig. 1.**
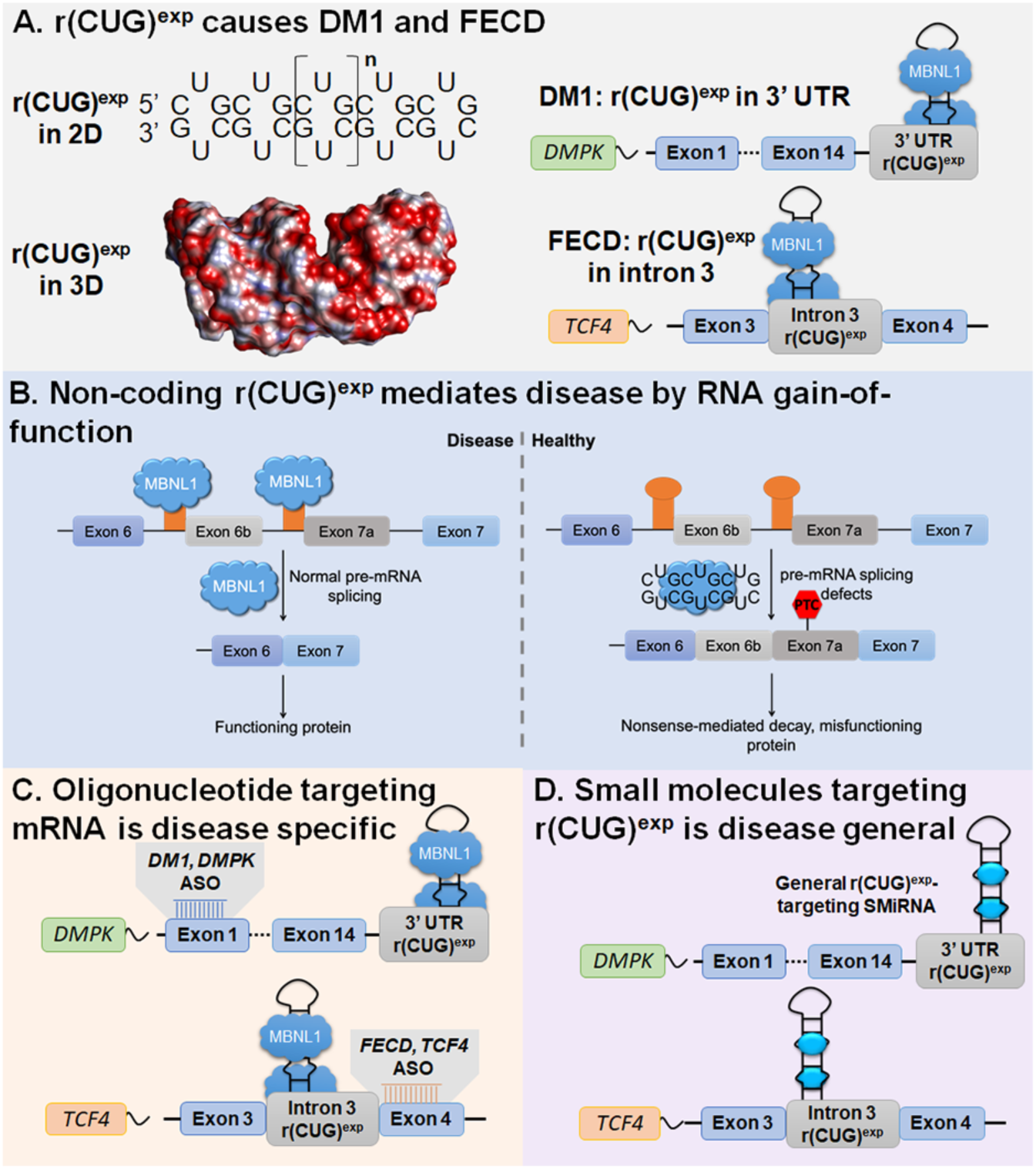
Mechanisms by which r(CUG)^exp^ causes DM1 and FECD and approaches to target the disease-causing RNAs with small molecules. (A) DM1 and FECD are caused by r(CUG)^exp^ found in the 3’ UTR of *DMPK* mRNA or in intron 3 of *TCF4* mRNA, respectively. The repeat expansion forms a structure containing repeating 1×1 UU internal loops that sequesters regulatory proteins such as MBNL1. (B) r(CUG)^exp^ causes disease via RNA gain-of-function in which it sequesters MBNL1 in nuclear foci, resulting in pre-mRNA splicing defects in genes that are regulated by MBNL1; the muscle specific chloride ion channel pre-mRNA (*CLCN1*) is shown as an example. (C) Using antisense oligonucleotides (ASOs) to target disease-causing RNAs requires customization for each transcript. (D) Developing small molecules that bind r(CUG)^exp^ structure offers a general way to target multiple diseases with the same compound.

Interestingly, it was recently discovered that r(CUG)^exp^ causes another disease, FECD, which manifests itself differently than DM1 as the repeat expansion is harbored in a different gene and in an intron. FECD is a dominantly inherited corneal disease that affects as many as 5% of Caucasian males, resulting in vision impairment (8). The FECD-causing r(CUG)^exp^ resides in intron 3 of the *Transcription Factor 4* (*TCF4*) pre-mRNA (4). Akin to DM1, r(CUG)^exp^ also sequesters MBNL1 in nuclear foci in FECD, causing pre-mRNA splicing defects (9-11). Recent studies have indicated that some *TCF4* mRNAs in FECD cells retain intron 3, a phenomenon that also occurs in other genetic disorders caused by intronic repeat expansions (12).

A common way to rescue the biology of microsatellite diseases is by the binding of antisense oligonucleotides (ASOs) that target unstructured regions in the coding mRNA (Fig. 1C) (13). Although targeting the repeat sequence itself has been investigated (14), specificity has been limited as tandem repeats are ubiquitous in the genome. Repeat-targeting ASOs also runs the risk of parallel knockdown of transcripts from the wild-type allele (15), which may give rise to loss-of-function phenotypes. Accordingly, an ASO is customized for each disease, even if two (or more) diseases are caused by the same repeat expansion.

An alternate approach to the sequence-based recognition of ASOs is the structure-based recognition of small molecules interacting with RNAs (SMIRNAs). That is, an SMIRNA would recognize the structure formed by the repeat expansions, for example the repeating array of 1 × 1 nucleotide U/U internal loops formed by r(CUG)^exp^ (16,17). Indeed, many small molecules have been developed to target the structure of r(CUG)^exp^ and improve DM1-associated defects in patient-derived cells and *in vivo* (16-20). Using a small molecule that directly recognizes the structure of RNA repeat expansions does not require customization across different diseases involving the same repeat motif, such as DM1 and FECD (Fig. 1D).

Herein, we describe the design of a small molecule that binds r(CUG)^exp^ with nanomolar affinity and is broadly selective. Indeed, the molecule directly engages the target, as studied by the target profiling method competitive chemical-cross linking and isolation by pull down (C-Chem-CLIP). As a result of this binding, we find that the compound improves r(CUG)^exp^-mediated defects similarly in both DM1 and FECD. In addition, the small molecule reduces the frequency of intron retention in *TCF4* in FECD-affected cells. The excised intron is subsequently degraded by hRRP6 and the nuclear exosome complex. Thus, RNA structure-specific ligands can be applied across diseases that are mediated by the same toxic structure. The impact of these compounds on metabolism of the RNA target can vary, depending on where the repeat expansion lies in the host gene, and in some cases may accelerate decay of the toxic element via quality control machinery.

## Results and Discussion

### Compound design, *in vitro* evaluation, and structural analysis

The most avid and bioactive ligands that have been designed to target repeat expansions of r(CUG)^exp^ have been dimers that contain two 5’C<u>U</u>G/3’G<u>U</u>C-binding modules (**H**) (16). A considerable challenge in targeting RNA structures, however, is developing low molecular weight, monomeric compounds that bind with high affinity and specificity. Thus, in this study, monomeric ligands that target r(CUG)^exp^ and affect biology broadly were developed. Previously, we reported that aryl diamines separated by a *bis*-benzimidazole scaffold was a favored sub-structure that targets r(CUG)^exp^, revealed by chemical similarity searching using **H** as query molecule (17,21). Compounds that were found to bind r(CUG)^exp^ and inhibit formation of the r(CUG)^exp^-MBNL1 complex include **1, 2**, and **3** (Fig. 2A) (17,21). Further, modeling studies of **1** (**H1**) suggested that the amino side chains interacted, in part, with the phosphodiester backbone (21).

**Fig. 2.**
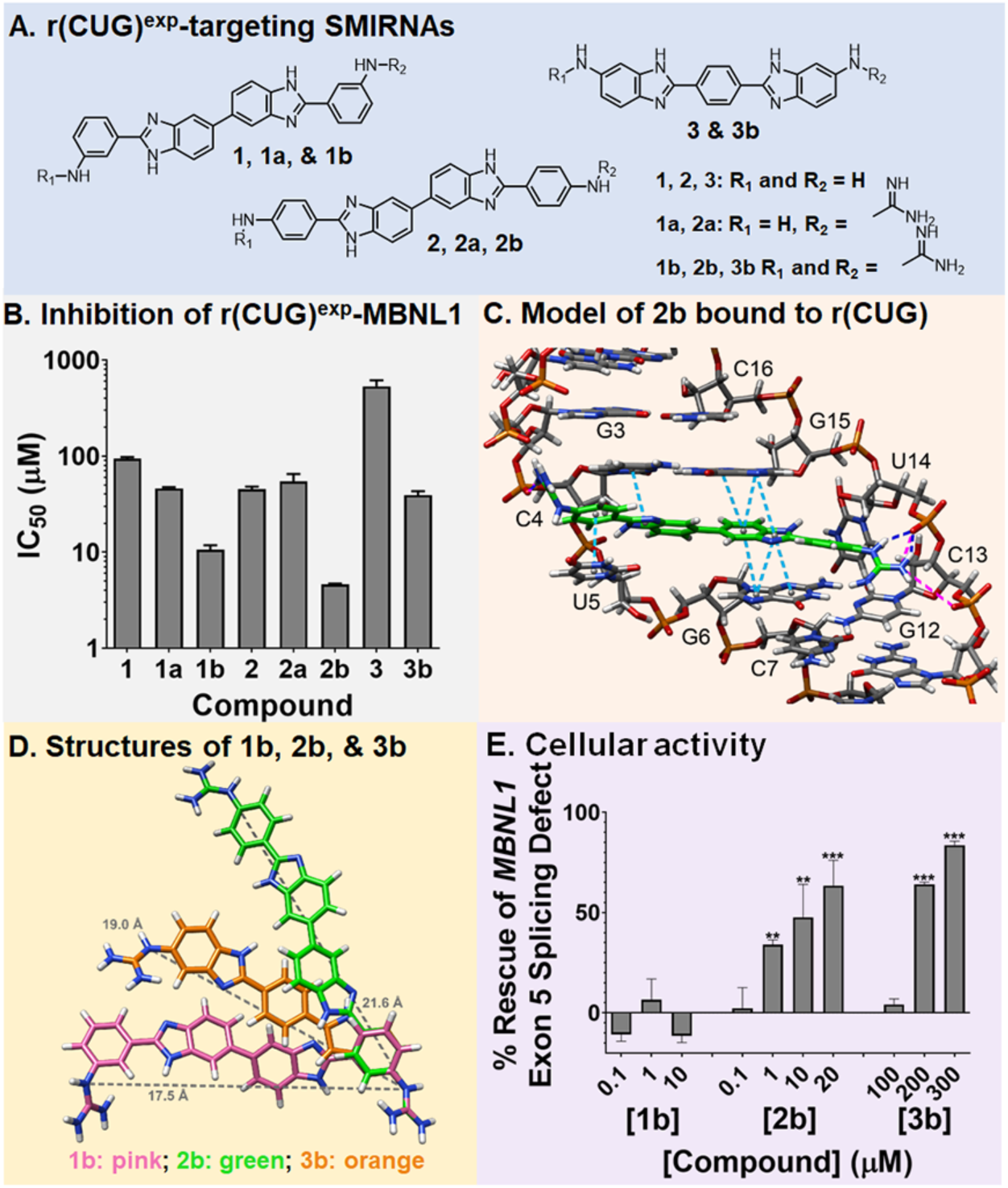
Design of compounds that bind r(CUG)^exp^. (A) Structures of small molecules interacting with RNA (SMIRNAs) that target r(CUG)^exp^. (B) Studying compounds *in vitro* via a previously reported TR-FRET assay to identify compounds that potently inhibit the r(CUG)^exp^-MBNL1 complex. (C) 3D model of interactions of **2b** with the loops formed by r(CUG)^exp^. Pink lines are ionic interactions, dark blue lines are hydrogen bonding interactions, and light blue lines are stacking interactions. (D) Structures of **1b** (pink), **2b** (green), and **3b** (orange) used to calculate the distance between guanidine substituents (gray line). (D) Rescue of *MBNL1* exon 5 splicing DM1 fibroblasts by **1b, 2b**, and **3b** (n = 3). **, *P* < 0.01; ***, *P* < 0.001, as determined by a one-way ANOVA by comparison to untreated cells. Error bars indicate SD.

One way to enhance activity of RNA-targeting ligands is through guanidinylation, which increases and distributes charge such that interactions are formed with the phosphodiester backbone. Thus, lead compounds **1, 2**, and **3**, were treated with cyanamide under acidic conditions to obtain singly (**1a** and **2a**) and doubly guanidinylated (**1b, 2b, 3b**) molecules in one pot reactions (Fig. 2A & Fig. S1). Each compound was evaluated *in vitro* for inhibition of the r(CUG)_12_-MBNL1 complex using a previously developed time-resolved fluorescence resonance energy transfer (TR-FRET) assay (Fig. 2B) (22). Relative to the parent compounds, each derivative provided a more potent *in vitro* response, and potency increased with the number of guanidine substituents. In particular, the doubly guanidinylated compounds were at least 10-fold more potent than the parent molecules. The affinity of the most potent compound, **2b** (IC_50_ = 4.6±0.1 µM), which is also the most potent *in cellulis* (*vide infra*), was measured by a direct binding assay. The compound bound avidly to a model of r(CUG)^exp^ with a K_d_ of 40±6 nM, while there was no measurable binding to a full base-paired RNA lacking 1×1 UU internal loops, an AT rich DNA hairpin, or other RNA repeat expansions including models of the c9ALS/FTD RNA [r(G_4_C_2_)^exp^] and the HD RNA [r(CAG)^exp^] (K_d_ > 5000 nM) (Fig. S2).

To study the features of molecular recognition, we completed dynamic docking to obtain a structural model of **2b** bound to an RNA with one copy of a 5’C<u>U</u>G/3’G<u>U</u>C internal loop motif (Fig. 2C and Fig. S3). Indeed, multiple interactions drive molecular recognition of the target. Previous biophysical studies have shown that the 5’C<u>U</u>G/3’G<u>U</u>C loops in the DM1 RNA are dynamic, adopting conformations in which the Us form 0, 1 (most populated), and 2 hydrogen bonds (23). Upon binding, **2b** changes the structure of the RNA such that the loop nucleotides no longer stack on the closing base pairs, replaced by the stacking of **2b**. The compound spans the major groove to form inter-strand interactions with the phosphodiester backbone and an array of hydrogen bonds between its *bis*-benzimidazole imino protons and the nucleobases and the sugar. Collectively, these modeling studies support that specific interactions and shape complementarity drive complex formation between **2b** and r(CUG) repeats.

To gain further insights into the shape complementarity and its role in driving complex formation, we overlaid the structures of **1b, 2b**, and **3b**, which may provide a rationale for **2b**’s enhanced potency (Fig. 2D). It is apparent that both the size of **2b** and the positioning of its functional groups affords the most complementarity to the structure formed by r(CUG)^exp^. For example, the shape presented by the 1×1 nucleotide UU internal loops is best spanned by **2b** such that the two ends of the ligand can interact with two phosphates. In contrast, **1b** does not have its guanidinium groups in the proper orientations to bind both phosphates simultaneously while compound **3b** is too short to accommodate both of these interactions simultaneously.

### Designer ligands improve various DM1 cellular defects

The doubly guanidinylated compounds (**1b, 2b**, and **3b**), the most potent for inhibition of the r(CUG)^exp^-MBNl1 complex *in vitro*, were studied for improving various DM1 defects in patient-derived fibroblasts at concentrations where no toxicity was observed (Fig. S4). Compounds were first tested for their ability to rescue the *MBNL1* exon 5 splicing defect, as MBNL1 protein regulates the splicing of its own pre-mRNA, in DM1 patient-derived fibroblasts; that is, *MBNL1* exon 5 alternative splicing is dysregulated in DM1 fibroblasts (24). While **1b** failed to improve the splicing pattern, **2b** and **3b** both rescued splicing dose-dependently, with statistically significant improvement observed at 1 µM and 200 µM, respectively (Fig. 2E and Fig. S5). Notably, **2b** did not affect *MBNL1* exon 5 inclusion in wild-type fibroblasts (Fig. S6) nor a NOVA-regulated splicing event in DM1 fibroblasts (Fig. S7). **2b** also reduced the number of r(CUG)^exp^- and MBNL1-containing nuclear foci in DM1 fibroblasts, as determined by fluorescence *in situ* hybridization (FISH) and MBNL1 immunofluorescence (Fig. S8). The observed improvements of DM1-associated molecular defects were not due to transcriptional inhibition of *DMPK* levels, which were unaffected by **2b** treatment as determined by RT-qPCR (Fig. S9).

As **2b** was clearly the most potent, we studied it further in DM1patient-derived myotubes, differentiated from fibroblasts (25). In agreement with studies in fibroblasts, **2b** reduced the number of nuclear foci/cell at 10 µM dose (Figs. 3A and 3B) and improved the *MBNL1* exon 5 splicing defect dose-dependently, significantly upon treatment with 1 µM compound (Fig. 3C and Fig. S10). As expected from studies in fibroblasts, **2b** did not affect *MBNL1* exon 5 inclusion in wild-type fibroblasts or myotubes (Fig. 3D and Fig. S6), and no effect was observed on NOVA-regulated *MAP4K4* exon 22a splicing (26) (Fig. 3E and S7). Additionally, the levels of *DMPK* mRNA were not affected by **2b** treatment, as measured by RT-qPCR (Fig. 3F), indicating that **2b** binds the RNA rather than inhibits transcription. Collectively, these studies in both DM1 patient-derived fibroblasts and myotubes indicate that **2b** acts in a manner that is consistent with selectively improving DM1 disease biology.

**Fig. 3.**
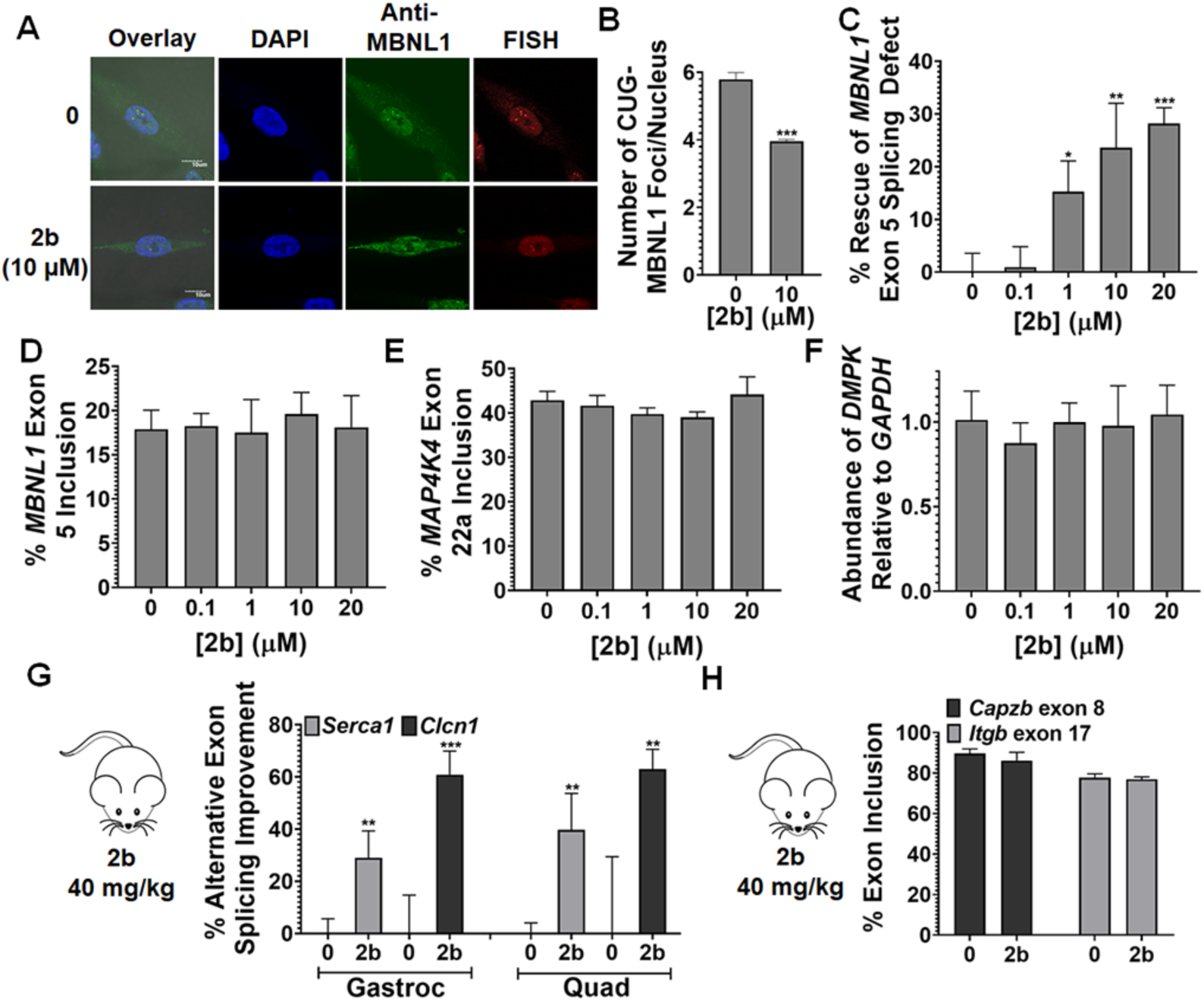
Compound **2b** alleviated molecular defects in DM1 myotubes and a DM1 mouse model. (A) Representative images of r(CUG)^exp^-MBNL1 foci in DM1 myotubes, imaged by MBNL1 immunostaining and RNA fluorescence *in situ* hybridization (FISH). (B) Quantification of the number of r(CUG)^exp^-MBNL1 foci/nucleus in DM1 myotubes (n = 3; 40 nuclei counted/replicate). *** *P* < 0.001, as determined by a two-tailed Student *t*-test by comparison to untreated cells. (C) Ability of **2b** to improve *MBNL1* exon 5 splicing in DM1 myotubes (n = 3). * *P* < 0.05, ** *P* < 0.01, *** *P* < 0.001, as determined by one-way ANOVA by comparison to untreated cells. (D) Evaluation of *MBNL1* exon 5 splicing in wild-type myotubes (n = 3). (E) Evaluation of *MAP4K4* exon 22a splicing in DM1 myotubes (n = 3). (F) Evaluation of *DMPK* levels in DM1 myotubes treated with **2b**, as determined by RT-qPCR with gene-specific primers (n = 3). (G) Ability of **2b** to improve *Serca 1* exon 22 and *Clcn1* exon 7a splicing in *HSA*^LR^ mice. “Gastroc” indicates gastrocnemius and “Quad” indicates quadriceps (n = 4 mice/group). **, *P* < 0.01; *** *P* < 0.001, as determined by a two-tailed Student *t*-test relative to vehicle treated (0). (H) Evaluation of *Capzb* exon 8 and *Itgb* exon 17 splicing (non-MBNL1 regulated events) in *HSA*^LR^ mice treated with **2b** (n = 4 mice/group). Error bars represent SD for all panels.

### Designer ligands directly bind to r(CUG)^exp^ in DM1 patient-derived cells

We next studied if the different potencies of **2b** and **3b** could be traced to the degree of r(CUG)^exp^ target engagement in cells. To measure target occupancy, we utilized the method chemical-cross linking and isolation by pull down (Chem-CLIP) (27). In Chem-CLIP, a chemical probe is appended with cross-linking (chlorambucil, CA) and purification (biotin) modules (Fig. 4A) (27). The Chem-CLIP probe undergoes a proximity-based cross-linking reaction to conjugate the structure-targeting ligand to the RNAs that it binds in cells. The cross-linked RNA can then be purified using streptavidin. Chem-CLIP probe **2H-K4NMeS-CA-Biotin** (16) binds r(CUG)^exp^ and pulls down mutant, but not wild-type, *DMPK* mRNA in DM1 fibroblasts, enriching the RNA by 20-fold (100 nM, Fig. 4A and Fig. S11).

**Fig. 4.**
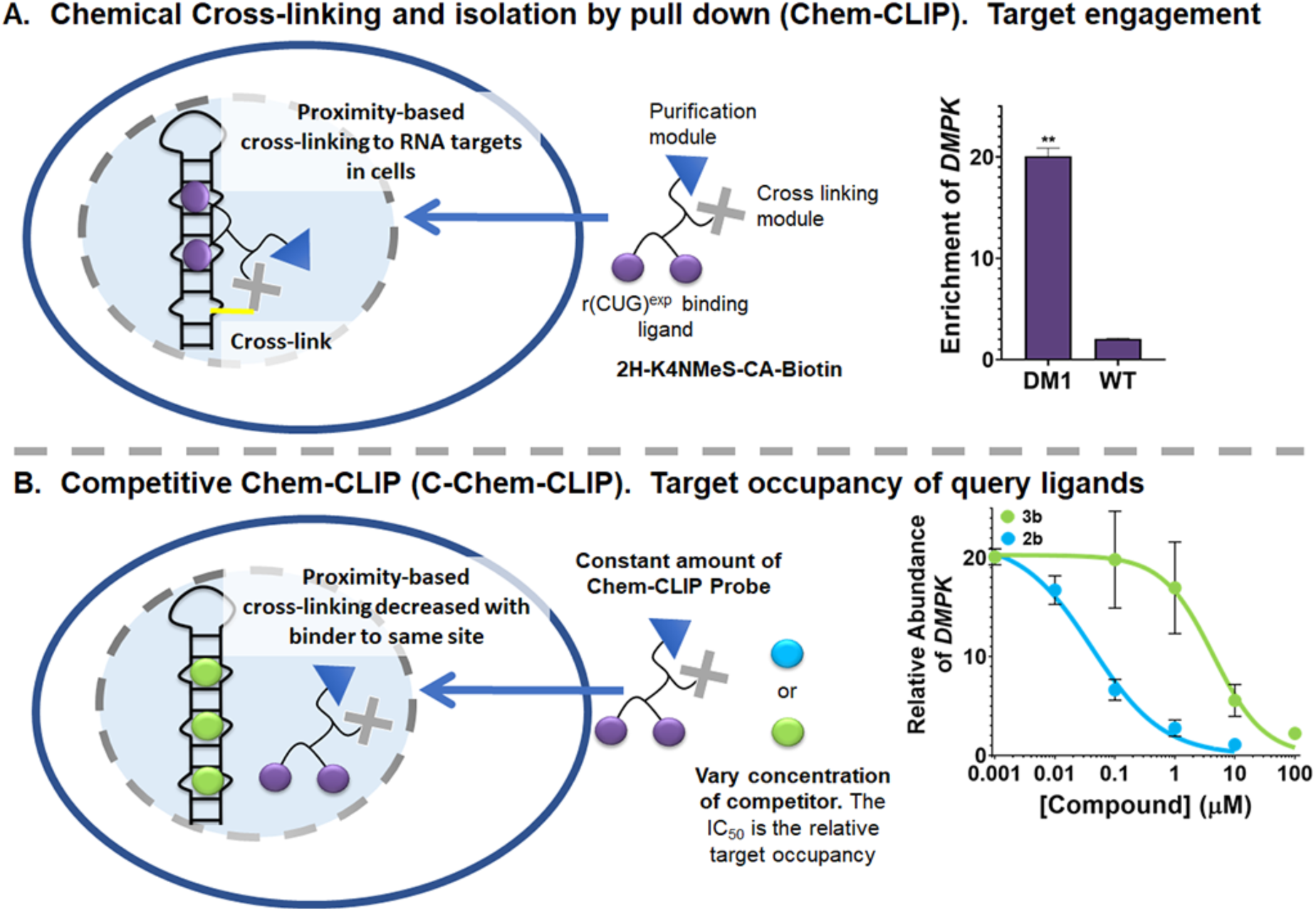
Compounds designed to target r(CUG)^exp^ engage the target in DM1 fibroblasts, as determined by C-Chem-CLIP. (A) Chem-CLIP target engagement in cells using Chem-CLIP probe **2H-K4NMeS-CA-Biotin**, which selectively cross-links and enriches r(CUG)^exp^-containing *DMPK* in DM1 fibroblasts. The probe does not enrich *DMPK* in wild-type fibroblasts. (n = 3 for both DM1 and WT fibroblasts). Data for WT fibroblasts was previously collected in Rzuczek et al. (16). **, *P* < 0.01, as determined by a two-tailed Student *t*-test. Error bars represent SD. (B) Competitive Chem-CLIP (C-Chem-CLIP) to study target engagement by **2b** and **3b**. DM1 fibroblasts were co-treated with 100 nM of **2H-K4NMeS-CA-Biotin** and varying concentrations of **2b** or **3b** to calculate the IC_50_s, or relative target occupancy in cells (n = 3). For **2b** the IC_50_ is 38±9 nM, and for **3b** the IC_50_ is 3400±60 nM. Error bars indicate SD.

Thus, this Chem-CLIP probe is suitable to study if ligands bind to r(CUG)^exp^ in cells by using competition experiment, or C-Chem-CLIP. In C-Chem-CLIP, cells are co-treated with the molecule of interest, in this case **2b** and **3b**, at varying concentrations in the presence of a constant concentration of the Chem-CLIP probe, in this case **2H-K4NMeS-CA-Biotin** (100 nM; Fig. 4B). Occupancy of the query ligand is calculated by its ability to inhibit the pull down of mutant DMPK mRNA. In these studies, **2b** had an IC_50_ of 38±9 nM, while **3b** had an IC_50_ of 3,400±60 nM, a difference of 100-fold for modest changes in compound structure (Fig. 4B). Since many molecules of MBNL1 and many molecules of compound can each bind r(CUG)^exp^, target occupancy can account for differences in cellular potency, at least in part. Interestingly, this difference in relative cellular occupancy in C-Chem-CLIP is greater than the relative difference in the ability of the two ligands to inhibit the formation of the r(CUG)^exp^-MBNL1 complex (Fig. 2B). This suggest that factors other than *in vitro* potency affect cellular activity and can include uptake and target selectivity that is read out in C-Chem-CLIP, which provides a holistic view at factors that affect RNA target occupancy in cells.

### Designer small molecule 2b rescues splicing *in vivo*

Lastly, we studied the therapeutic potential of **2b** in the *HSA*^LR^ mouse model of DM1 (28). This transgenic mouse model expresses a human skeletal actin (*HSA*) transgene containing r(CUG)_220_ and has several hallmarks of human disease including pre-mRNA splicing defects (28). To study if **2b** can improve these disease defects *in vivo, HSA*^LR^ mice were treated with 40 mg/kg of **2b** via intraperitoneal (*i.p.*) injection every day for 7 days. The compound was well tolerated, and analysis of RNA isolated from the gastrocnemius and quadriceps muscles after dosing showed specific improvement of two deregulated splicing events, *Serca1* exon 22 and *Clcn1* exon 7A (Fig. 3G and Fig. S12). These changes in splicing outcomes were specific to DM1, as two non-MBNL1 regulated splicing events, *Capzb* exon 8 and *Itgb* exon 17, were not affected (Fig. 3H and Fig. S13). Thus, **2b** potently and specifically improve DM1associated defects in patient-derived cell lines and *in vivo*.

### Designer small molecule 2b that targets r(CUG)^exp^ improved disease pathways in FECD

The corneal disease FECD is caused by r(CUG)^exp^ located in intron 3 of *TCF4* transcript (4). FECD causes thinning of the cornea that ultimately may require corneal transplantation. In contrast to DM1, it is not a rare disease, but rather a common disease that affects 5% of male Caucasians over the age of 40 (8). The disease is manifest at the molecular levels by similar biochemical events to that observed in DM1, *i.e.* r(CUG)^exp^ forms foci with MBNL1 and causes pre-mRNA splicing defects (9,10). Thus, we investigated whether **2b** also improves disease defects in FECD patient-derived corneal endothelium cells.

As determined by a cell viability assay, no measurable toxicity was observed up to 20 µM of **2b** (Fig. S14). We then studied whether **2b** could reduce r(CUG)^exp^-MBNL1 foci. As was observed in DM1, a reduction in the number of foci per cell was observed in FECD cells when treated with 10 µM of **2b**, from 3.6±0.2 to 2.7±0.3 (*P*<0.01; Fig. 5A and Fig. S15). As expected, **2b** also improved FECD-associated pre-mRNA splicing defects. A dose-response profile was observed for rescue of the *MBNL1* exon 5 splicing by **2b** (Fig. 5B and Fig. S15), similar to that observed in DM1 cells (Fig. 2E and 3C). In particular, statistically significant rescue was observed in low µM range in FECD cells (Fig. 5B). The compound had no effect on NOVA-regulated *MAP4K4* exon 22a splicing, and similar to wild-type fibroblast studies described above, no effect on *MBNL1* exon 5 splicing was observed in wild-type corneal endothelium cells, demonstrating the functional selectivity of **2b** in FECD cells (Fig. S15 and Fig. S16).

**Fig. 5.**
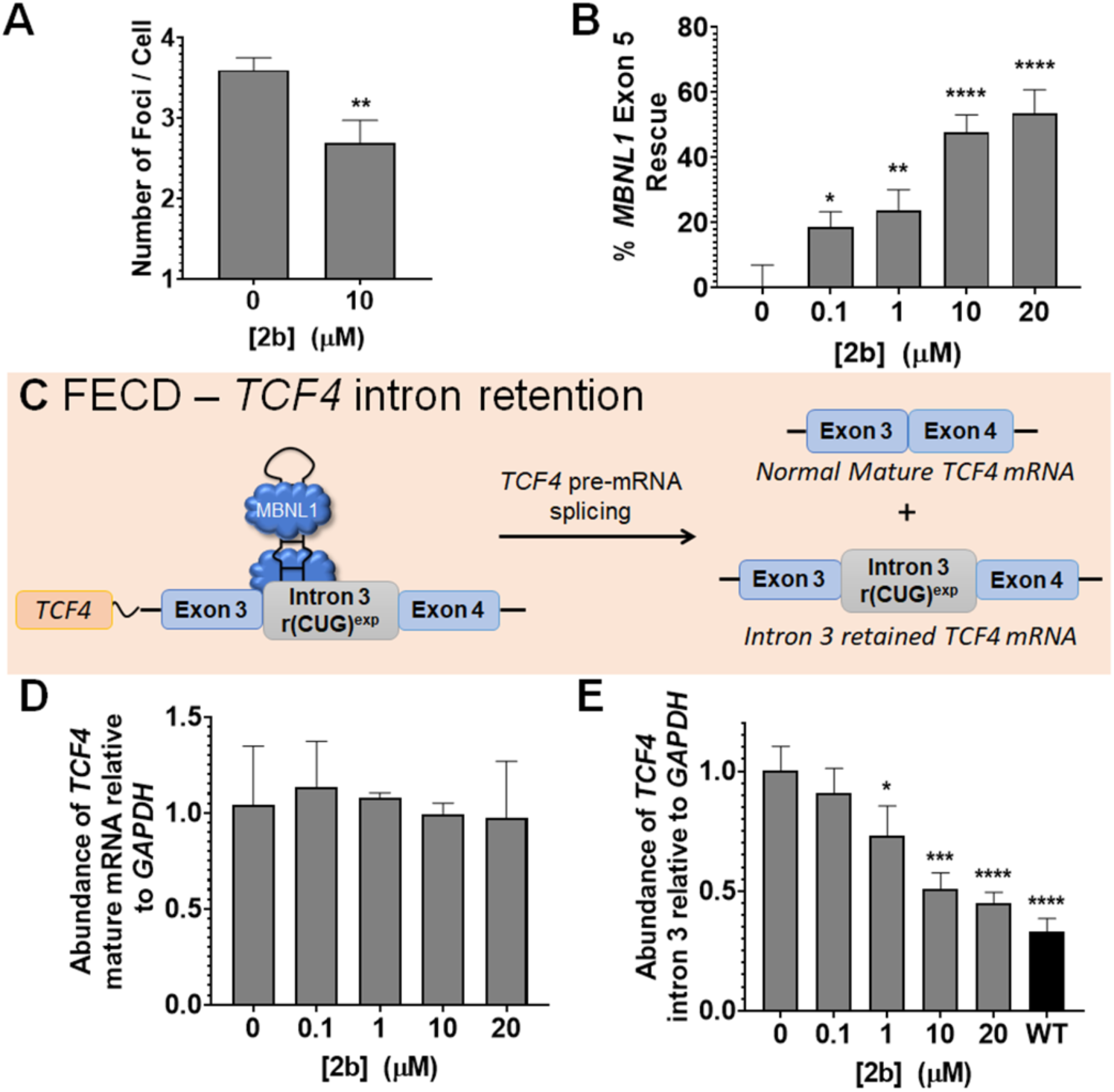
Compound **2b** alleviates molecular defects in FECD cells. (A) Quantification of the number of r(CUG)^exp^-MBNL1 foci/nucleus (n = 3, 40 nuclei counted/replicate). **, *P* < 0.01, as determined by a two-tailed Student t-test. (B) Effect of **2b** on *MBNL1* exon 5 pre-mRNA splicing (n = 3). *, *P* < 0.05; **, *P* < 0.01; ****, *P* < 0.0001, as determined by one-way ANOVA compared to untreated cells (“0”). Error bars represent SD. (C) Retention of *TCF4* intron 3 in FECD-affected cells. (D) Effect of **2b** on *TCF4* mature mRNA, as determined by RT-qPCR (n = 3). (E) Effect of **2b** on intron 3-containing *TCF4* levels in FECD cells, compared to WT levels, as determined by RT-qPCR (n = 3). *, *P* < 0.05; ***, *P* < 0.001; ****, *P* < 0.0001, as determined by one-way ANOVA compared to untreated cells (“0”).

### Designer small molecule 2b triggers decay of the intronic r(CUG) repeat expansion via the RNA exosome

Next, we studied the effect of **2b** on the levels of the *TCF4* transcript and intron 3, as previous studies have shown that intronic RNA repeat expansions, particularly those that are GC-rich, cause intron retention in the mature mRNAs (Fig. 5C) (12). In myotonic dystrophy type 2 (DM2), intron retention is due, at least in part, to formation of the r(CCUG)^exp^-MBNL1 complex (29). Inhibition of this complex via small molecule treatment triggered decay of the retained intron, but there was no mechanistic basis for this observation, for example which enzymes were responsible for the intron’s degradation (29). To study the levels of *TCF4* in FECD cells, we performed RT-qPCR using primers for mature *TCF4* mRNA and primers specific for r(CUG)^exp^-containing intron 3. The levels of mature *TCF4* mRNA in FECD cells were not affected upon compound treatment (Fig. 5D). The levels of transcripts containing intron 3, however, were decreased in a dose-dependent manner (Fig. 5E). In particular, as little as 1 µM of **2b** reduced levels of the retained intron, and reversion to levels observed in wild-type cells was nearly achieved at 20 µM (Fig. 5E). Importantly, transcripts containing intron 3 were not affected in wild-type cells (Fig. S16).

Next, we investigated the mechanism of decreased intron 3-retained *TCF4* transcripts. Following removal by the spliceosome, introns are processed by debranching enzymes and then degraded by exonucleases. We used a siRNA approach to ablate various exonucleases that may be contributing to the decay of the r(CUG)^exp^-containing intron. After siRNA knockdown, FECD cells were then treated with **2b**; if the exonuclease is contributing to degradation of the intron, **2b** would therefore lose its ability to reduce intron 3-containing *TCF4* transcripts. We focused our studies on two different complexes responsible for RNA decay, 5’-3’ Exoribonuclease 1 (XRN1) and hRRP6, a catalytic domain of the exosome [also known as Polymyositis/Scleroderma Autoantigen 100 KDa (PM/Scl100)] (30). The human exosome contains 11 subunits including two catalytic domains, hRRP6 and hRRP44 (31).

When *XRN1* expression was knocked down, no significant change in **2b**’s activity to reduce *TCF4* intron 3 levels was observed, suggesting that XRN1 is not involved in the degradation of the intron (Fig. 6A and Fig. S17). In contrast, siRNA knock down of hRRP6 reduced the ability of **2b** to decrease the levels of intronic r(CUG)^exp^. Co-treatment of **2b** and a control siRNA did not affect the reduction of intron 3 by **2b** (Fig. 6B and Fig. S17). Collectively, these results strongly suggest that the native decay of the retained intron triggered by **2b** is in part regulated by the multi-protein intracellular exosome complex through the hRRP6 exonuclease (Fig. 6C).

**Fig. 6.**
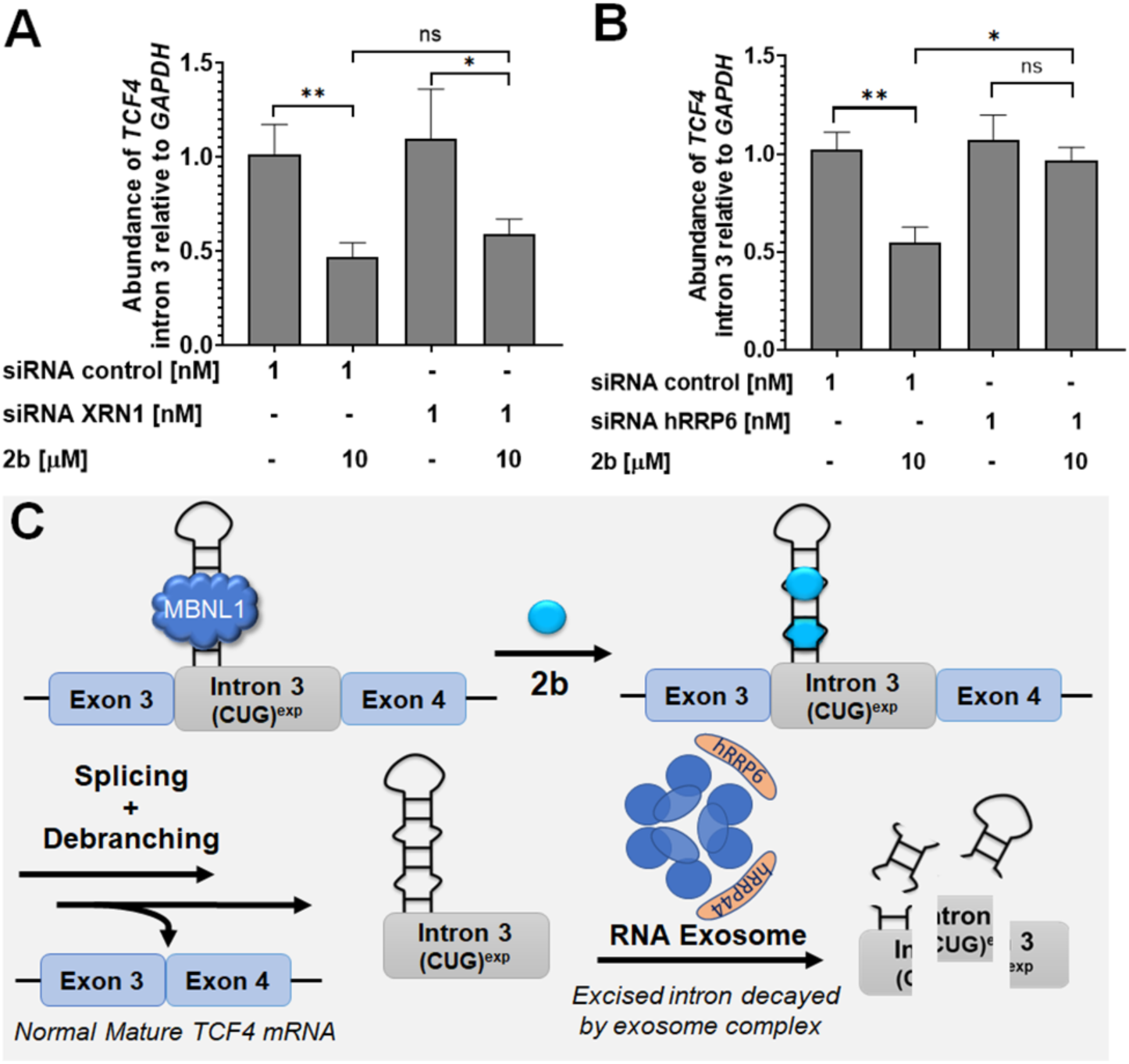
Mechanism of r(CUG)^exp^-containing intron 3 decay in FECD cells. (A) Effect of the siRNA targeting *XRN1* in combination with **2b** on *TCF4* intron 3 levels, as determined by RT-qPCR. Knock-down of *XRN1* had no effect on *TCF4* intron 3 decay induced by **2b** (n = 3). \**P*, < 0.05; \*\**P*, < 0.01, as determined by a two-tailed Student t-test. (B) Effect of the siRNA targeting *hRRP6* in combination with **2b** on *TCF4* intron 3 levels, as determined by RT-qPCR (n = 3). Knock-down of *hRRP6*, the catalytic domain of the exosome complex, decreases *TCF4* intron 3 decay generated by **2b**. \**P*, < 0.05; \*\**P*, < 0.01, as determined by a two-tailed Student t-test. Error bars represent SD. (*C*) Proposed decay mechanism, where the exosome complex plays a key role in the decay pathway. Exosome nucleolytic domains are represented in orange including the hRRP6 subunit.

### Implications on the generality of ligands that target RNA structure

Oligonucleotides, such as, ASOs or siRNAs, are increasingly being used to treat genetic diseases. Their sequence-based recognition is biased for regions in an RNA target that lack structure (32). Because repeating sequences are found throughout the genome, ASOs that target them lack selectivity, either through decay by an RNase H-dependent mechanism (18) or by steric blockade such as with morpholino oligonucleotides (14). Therefore, ASOs for microsatellite diseases have been developed that recognize sequences unique to each gene, rather than the repeat expansion itself. Thus, a single ASO could not be used for both FECD and DM1, despite sharing a common causative agent, r(CUG)^exp^.

Small molecule targeting of RNA is still in its relative infancy, and many more studies will be required to truly assess their potential to ameliorate disease pathways and thus serve as therapeutics or chemical probes. Repeat expansions can indeed be targeted in an allele-specific manner by using small molecules that target RNA structure (16). This selectivity can be traced at least in part to the differences in structure formed by long toxic repeats, such as r(CUG)^exp^, and short, non-pathological repeats, which do not generally form stable structures (18).

Herein, we show that ligands that recognize the toxic structure formed by r(CUG)^exp^ can be applied across diseases to study r(CUG)^exp^-mediated biology and with potential as treatments for all r(CUG)^exp^-mediated diseases, provided the compounds could be developed into clinical candidates. There are three known diseases mediated by r(CUG)^exp^, DM1, FECD, and Huntington’s-like 2 (HDL2) (33,34). Such an approach would be even more far-reaching for small molecules that bind r(CAG)^exp^ and deactivate disease pathways, as it operates in nearly a dozen different diseases (35). That is, the overall implications of our studies are that many different diseases caused by the same repeat expansion could be treated with a single ligand and that small molecule mechanisms to alleviate disease biology can be unique to each disease depending on where the repeat expansion is harbored (UTR vs. intron, for example).

### Targeted degradation is a novel mode of action for small molecules that target RNA structures

There is an intense interest in targeted degradation as a strategy to ameliorate disease states. Various modalities have been deployed to target the degradation of biomolecules, including proteolysis targeting chimeras (PROTACs) for proteins (36) and ASOs and ribonuclease targeting chimeras (RIBOTACs) (37) for RNAs. All three target biomolecules for destruction by using ligands that possess larger molecular weight and fall outside of Lipinksi guidelines (38); for PROTACs and RIBOTACs, this is because of their chimeric nature to bind the target and recruit an enzyme.

The mode of action of **2b** suggests that intrinsic RNA decay pathways can be harnessed to accelerate the degradation of a disease-causing RNA. Thus, targeted degradation functionality can be accessed by using simple binding ligands with low molecular weights that may be more within the purview of drug-like chemical space. There is a myriad of RNA quality control pathways that can process RNAs in many different manners. It is very likely that specific targeting of RNA structure can facilitate the recognition of RNA by the quality control machinery by using very simple ligands such as **2b**. Furthermore, the observed selectivity of a small molecule could be even greater than expected if off-target binding occurs in a non-functional site that has no effect on downstream biology (39). Thus, although a binding site may not be unique, affecting function and transcript abundance can be selective because only certain transcripts, such as retained repeats in introns, are sensitive to binding of ligands.

### Conclusions and Outlook

Although targeting RNA structures with small molecules are still at an early stage, the utility of such structure-binding ligands as chemical probes has been established. However, the clinical development of such compounds is only beginning to emerge. Small molecules can modulate RNA function in various ways, and the mechanistic underpinnings in this study show that one way is through stimulation of exosomal decay. Perhaps, small molecules can stimulate other RNA quality control pathways. Several other diseases are mediated by repeat expansions in retained introns (12) including as DM2 (29) and c9ALS/FTD (40), and as such it will be interesting to see if the same pathway can be activated with ligands that target these repeat expansions. If so, these targets may be more druggable than they immediately appear.

## Supporting information

Full supporting information

## Acknowledgements

We thank J. Childs-Disney for experimental advice and Prof. Denis Furling [Centre de Recherche en Myologie (UPMC/Inserm/CNRS), Institut de Myologie], for his generous gift of myotube cell lines used in this paper. We also thank the agencies that funded this work including the National Institutes of Health (DP1-NS096898 to M.D.D., F31 NS110269 to A.J.A., and P50-NS048843 to C.A.T.), the Muscular Dystrophy Association (grant 380467 to M.D.D.), the Myotonic US Fellowship Research grant (to R.I.B. and S.C.) and to the National Ataxia Foundation fellowship research grant (to R.I.B.). We would also like to

## Author Contributions

M.D.D. directed the study, conceived of ideas, and designed experiments. A.J.A., R.I.B., S.G.R. designed experiments, synthesized compounds, and conducted biochemical and cellular studies. S.C., J.L.C, K.W.W, and I.Y. performed modeling studies. Z.T. and C.A.T. performed experiments in mice. M.P. and A.S.J. provided cell lines.

## References

1. The Huntington’s Disease Collaborative Research Group, A novel gene containing a trinucleotide repeat that is expanded and unstable on Huntington’s disease chromosomes. Cell 72, 971–983 (1993).

2. M. DeJesus-Hernandez et al., Expanded GGGGCC hexanucleotide repeat in noncoding region of C9ORF72 causes chromosome 9p-linked FTD and ALS. Neuron 72, 245–256 (2011).

3. J. D. Brook et al., Molecular basis of myotonic dystrophy: Expansion of a trinucleotide (CTG) repeat at the 3′ end of a transcript encoding a protein kinase family member. Cell 68, 799–808 (1992).

4. E. D. Wieben et al., A common trinucleotide repeat expansion within the transcription factor 4 (TCF4, E2-2) gene predicts Fuchs corneal dystrophy. PLoS One 7, e49083–e49083 (2012).

5. K. L. Taneja, M. McCurrach, M. Schalling, D. Housman, R. H. Singer, Foci of trinucleotide repeat transcripts in nuclei of myotonic dystrophy cells and tissues. J. Cell Biol. 128, 995–1002 (1995).

6. H. Jiang, A. Mankodi, M. S. Swanson, R. T. Moxley, C. A. Thornton, Myotonic dystrophy type 1 is associated with nuclear foci of mutant RNA, sequestration of muscleblind proteins and deregulated alternative splicing in neurons. Hum. Mol. Genet. 13, 3079–3088 (2004).

7. M. Nakamori et al., Splicing biomarkers of disease severity in myotonic dystrophy. Ann. Neurol. 74, 862–872 (2013).

8. D. W. Lorenzetti, M. H. Uotila, N. Parikh, H. E. Kaufman, Central cornea guttata. Incidence in the general population. Am. J. Ophthalmol. 64, 1155–1158 (1967).

9. J. Du et al., RNA toxicity and missplicing in the common eye disease Fuchs endothelial corneal dystrophy. J. Biol. Chem. 290, 5979–5990 (2015).

10. V. V. Mootha et al., TCF4 triplet repeat expansion and nuclear RNA foci in Fuchs’ endothelial corneal dystrophy. Invest. Ophthalmol. Vis. Sci. 56, 2003–2011 (2015).

11. M. A. Hale, N. E. Johnson, J. A. Berglund, Repeat-associated RNA structure and aberrant splicing. BBA Gene Regul. Mech. 1862, 194405 (2019).

12. Ł. J. Sznajder et al., Intron retention induced by microsatellite expansions as a disease biomarker. Proc. Natl. Acad. Sci. 115, 4234–4239 (2018).

13. M. D. Disney, Targeting RNA with small molecules to capture opportunities at the Intersection of chemistry, biology, and medicine. J. Am. Chem. Soc. 141, 6776–6790 (2019).

14. T. M. Wheeler et al., Reversal of RNA dominance by displacement of protein sequestered on triplet repeat RNA. Science 325, 336–339 (2009).

15. D. Jauvin et al., Targeting DMPK with antisense oligonucleotide improves muscle strength in myotonic dystrophy type 1 mice. Mol. Ther. Nucleic Acids 7, 465–474 (2017).

16. S. G. Rzuczek et al., Precise small-molecule recognition of a toxic CUG RNA repeat expansion. Nat. Chem. Biol. 13, 188–193 (2017).

17. S. G. Rzuczek, M. R. Southern, M. D. Disney, Studying a drug-like, RNA-focused small molecule library identifies compounds that inhibit RNA toxicity in myotonic dystrophy. ACS Chem. Biol. 10, 2706–2715 (2015).

18. A. J. Angelbello et al., Precise small-molecule cleavage of an r(CUG) repeat expansion in a myotonic dystrophy mouse model. Proc. Natl. Acad. Sci. 116, 7799–7804 (2019).

19. J. Li et al., A Dimeric 2,9-Diamino-1,10-phenanthroline derivative improves alternative splicing in myotonic dystrophy type 1 cell and mouse models. Chemistry 24, 18115–18122 (2018).

20. A. H. Jahromi et al., Developing bivalent ligands to target CUG triplet repeats, the causative agent of myotonic dystrophy type 1. J. Med. Chem. 56, 9471–9481 (2013).

21. R. Parkesh et al., Design of a bioactive small molecule that targets the myotonic dystrophy type 1 RNA via an RNA motif–ligand database and chemical similarity searching. J. Am. Chem. Soc. 134, 4731–4742 (2012).

22. C. Z. Chen et al., Two high-throughput screening assays for aberrant RNA–protein interactions in myotonic dystrophy type 1. Anal. Bioanal. Chem. 402, 1889–1898 (2012).

23. R. Parkesh, M. D. Disney, M. Fountain, NMR spectroscopy and molecular dynamics simulation of r(CCGCUGCGG)2 reveal a dynamic UU internal loop found in myotonic dystrophy type 1. Biochemistry 50, 599–601 (2011).

24. D. P. Gates, L. A. Coonrod, J. A. Berglund, Autoregulated splicing of muscleblind-like 1 (MBNL1) pre-mRNA. J. Biol. Chem. 286, 34224–34233 (2011).

25. L. Arandel et al., Immortalized human myotonic dystrophy muscle cell lines to assess therapeutic compounds. Dis. Model. Mech. 10, 487–497 (2017).

26. J. Ule et al., Nova regulates brain-specific splicing to shape the synapse. Nat. Genet. 37, 844–852 (2005).

27. L. Guan, M. D. Disney, Covalent small-molecule–RNA complex formation enables cellular profiling of small-molecule–RNA interactions. Angew. Chem. Int. Ed. 52, 10010–10013 (2013).

28. A. Mankodi et al., Myotonic dystrophy in transgenic mice expressing an expanded CUG repeat. Science 289, 1769–1773 (2000).

29. R. I. Benhamou, A. J. Angelbello, E. T. Wang, M. D. Disney, A toxic RNA catalyzes the cellular synthesis of its own inhibitor, shunting it to endogenous decay pathways. Cell Chem. Biol. 27, 223–231.e224 (2020).

30. N. L. Garneau, J. Wilusz, C. J. Wilusz, The highways and byways of mRNA decay. Nat. Rev. Mol. Cell Biol. 8, 113–126 (2007).

31. C. Kilchert, S. Wittmann, L. Vasiljeva, The regulation and functions of the nuclear RNA exosome complex. Nat. Rev. Mol. Cell Biol. 17, 227–239 (2016).

32. T. A. Vickers, J. R. Wyatt, S. M. Freier, Effects of RNA secondary structure on cellular antisense activity. Nucleic Acids Res. 28, 1340–1347 (2000).

33. R. L. Margolis, D. D. Rudnicki, Pathogenic insights from Huntington’s disease-like 2 and other Huntington’s disease genocopies. Curr. Opin. Neurol. 29, 743–748 (2016).

34. T. Ratovitski et al., Quantitative proteomic analysis reveals similarities between Huntington’s disease (HD) and Huntington’s disease-like 2 (HDL2) human brains. J. Prot. Res. 15, 3266–3283 (2016).

35. H. Paulson, Repeat expansion diseases. Handb. Clin. Neurol. 147, 105–123 (2018).

36. K. G. Coleman, C. M. Crews, Proteolysis-targeting chimeras: Harnessing the ubiquitin-proteasome system to induce degradation of specific target proteins. Ann. Rev. Cancer Bio. 2, 41–58 (2018).

37. M. G. Costales et al., Small-molecule targeted recruitment of a nuclease to cleave an oncogenic RNA in a mouse model of metastatic cancer. Proc. Natl. Acad. Sci. 117, 2406–2411 (2020).

38. C. A. Lipinski, F. Lombardo, B. W. Dominy, P. J. Feeney, Experimental and computational approaches to estimate solubility and permeability in drug discovery and development settings. Adv. Drug Deliv. Rev. 23, 3–25 (1997).

39. M. G. Costales et al., Small molecule inhibition of microRNA-210 reprograms an oncogenic hypoxic circuit. J. Am. Chem. Soc. 139, 3446–3455 (2017).

40. M. Niblock et al., Retention of hexanucleotide repeat-containing intron in C9orf72 mRNA: implications for the pathogenesis of ALS/FTD. Acta. Neuropathol. Commun. 4, 18–18 (2016).

