## Supplementary material for "A small molecule that binds an RNA repeat expansion stimulates its decay via the exosome complex": Full supporting information

Supporting Information Contains:

- Table S1
- Supporting Figures S1 – S17
- Methods
- Synthetic Methods and Characterization

**Table S1.** Sequences of primers used for RT-PCR and RT-qPCR.

| Gene | Forward Primer (5'-3') | Reverse Primer (5'-3') | Purpose |
|---|---|---|---|
| *MBNL1* | GCTGCCCAATACCAGGTCAAC | TGGTGGGAGAAATGCTGTATGC | RT-PCR |
| *MAP4K4* | CCTCATCCAGTGAGGAGTCG | ATCACAGGAAAATCCCACCA | RT-PCR |
| *DMPK* | CGTGCAAGCGCCCAG | CTCCACCAACTTACTGTTTCATCCT | qPCR |
| *GAPDH* | AAGGTGAAGGTCGGAGTCAA | AATGAAGGGGTCATTGATGG | qPCR |
| *Clcn1* | TGAAGGAATACCTCACACTCAAGG | CACGGAACACAAAGGCACTG | RT-PCR |
| *Serca1* | GCTCATGGTCCTCAAGATCTCAC | GGGTCAGTGCCTCAGCTTTG | RT-PCR |
| *Itgb* | CCTACTGGTCCCGACATCATC | CTTCGGATTGACCACAGTTGTC | RT-PCR |
| *Capzb* | GCACGCTGAATGAGATCTACTTTG | CCGGTTAGCGTGAAGCAGAG | RT-PCR |
| *TCF4 mature mRNA* | ACGATGAGGACCTGACAC | GTCTGGGGCTTGTCACTCTT | qPCR |
| *TCF4 intron 3* | GAGAGAGGGAGTGAAAGAGAGA | GGCAATGTCCATTTCCATCT | qPCR |
| *EXOSC10* | CTCTTTGGACCTCACGACTGCT | AAGAGGCTCGCCTGCTTCTGAA | qPCR |
| *XRN1* | CCAGCAAAGCAGTCGTGGAGAA | CCACGACTCTAGCTTCCTCAAG | qPCR |

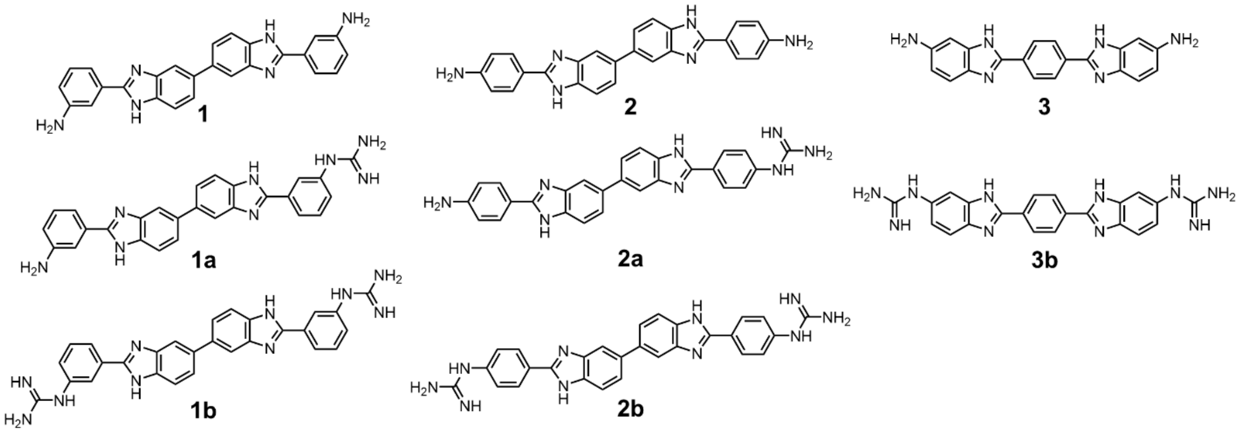

**Figure S1.** Chemical structures of all compounds tested.

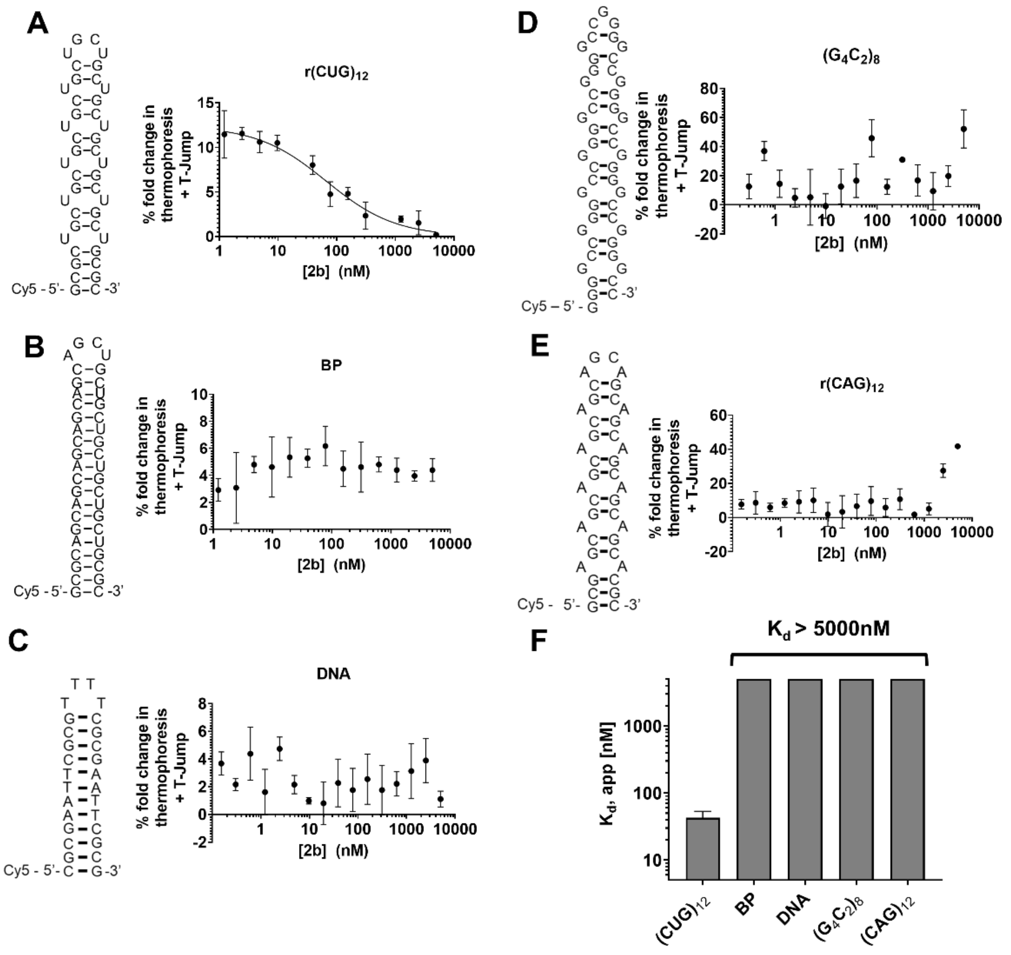


**Figure S2.** Compound **2b** binds avidly and selectivity to r(CUG)$_{12}$ vs. other RNA repeats, base-paired control RNA and AT-rich DNA, as measured by microscale thermophoresis (MST). (A) Thermophoresis analysis of **2b** bound to r(CUG)$_{12}$ (n = 3 independent experiments). (B) Thermophoresis analysis of **2b** with a base-paired control RNA (n = 3 independent experiments). (C) Thermophoresis analysis of **2b** with AT-rich (n = 3 independent experiments). (D) Thermophoresis analysis of **2b** with a r(G$_4$C$_2$)$_8$ (n = 3 independent experiments). (E) Thermophoresis analysis of **2b** with a r(CAG)$_{12}$ (n = 3 independent experiments). (F) Plot of the $K_{d,app}$ of **2b** and r(CUG)$_{12}$ ($K_d$ = 42 ± 6 nM) and to a base-paired control , AT-rich DNA, r(G$_4$C$_2$)$_8$ and r(CAG)$_{12}$ ($K_d$ > 5000 nM). Error bars indicate SD for all panels.

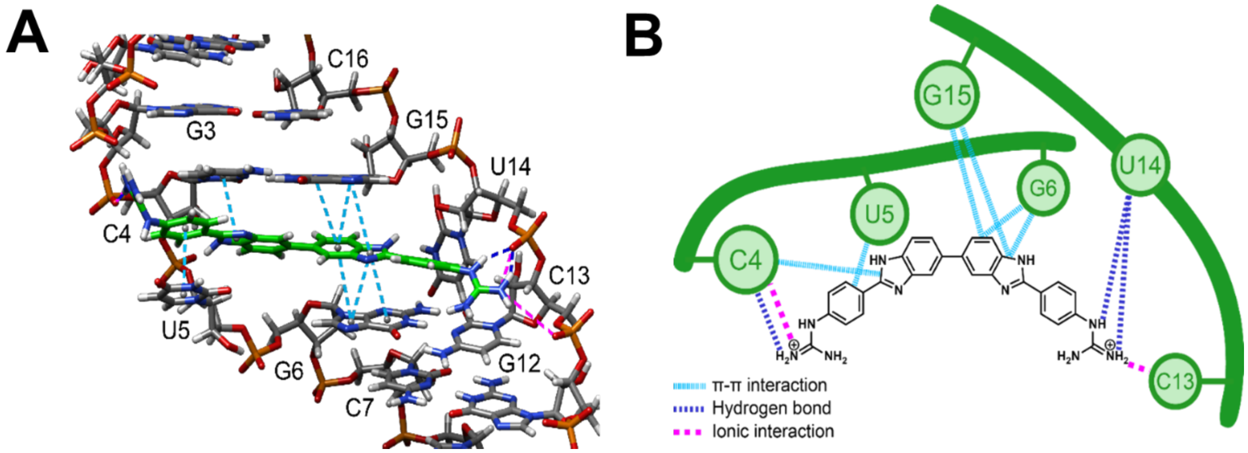


**Figure S3.** Model of **2b** bound to r(CCGC<u>U</u>GCGG)$_2$. (A) 3D model of interactions. Pink lines are ionic interactions, dark blue lines are hydrogen bonding interactions, and light blue lines are stacking interactions. (B) Schematic of interactions.

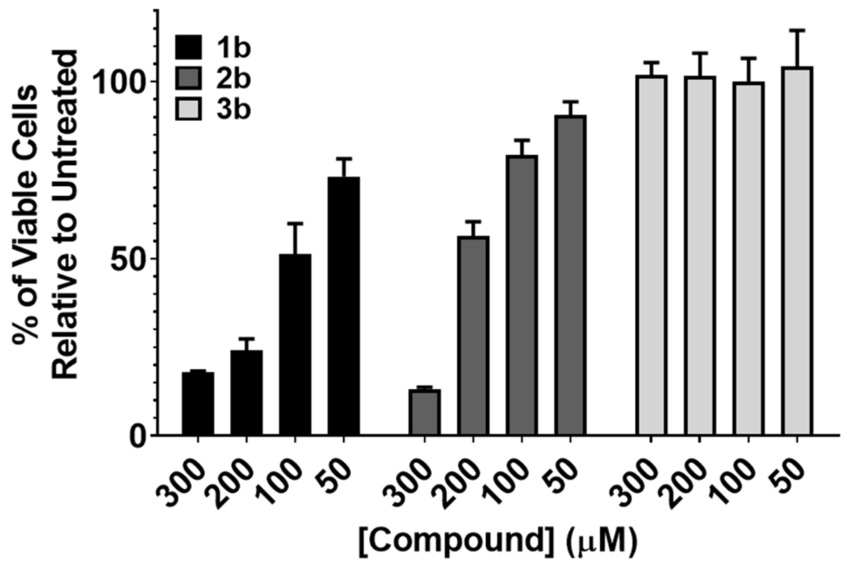


**Figure S4.** Toxicity of **1b**, **2b**, and **3b** in DM1 fibroblasts as determined by WST-1 cell viability reagent (n = 5). Error bars indicate SD.

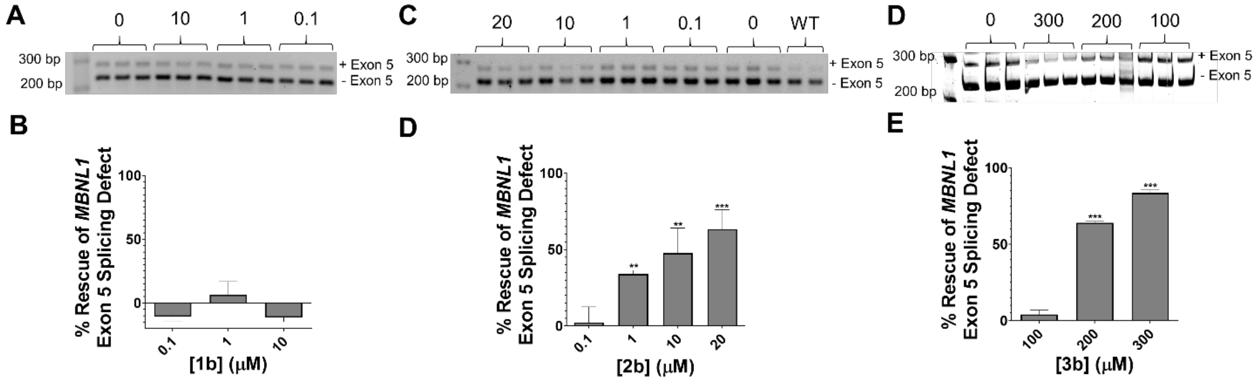


**Figure S5.** Rescue of the *MBNL1* exon 5 splicing defect by **1b**, **2b**, and **3b** in DM1 fibroblasts. (A) Representative gel image of *MBNL1* exon 5 splicing in DM1 fibroblasts treated with **1b**. (B) Quantification of *MBNL1* splicing rescue in DM1 fibroblasts treated with **1b**. (C) Representative gel image of *MBNL1* exon 5 splicing in DM1 fibroblasts treated with **2b**. (D) Quantification of *MBNL1* splicing rescue in DM1 fibroblasts treated with **2b**. (E) Representative gel image of *MBNL1* exon 5 splicing in DM1 fibroblasts treated with **3b**. (F) Quantification of *MBNL1* splicing rescue in DM1 fibroblasts treated with **3b**. For all panels, n = 3 and error bars indicate SD. **, $P < 0.01$; ***, $P < 0.001$, as determined by a one-way ANOVA.

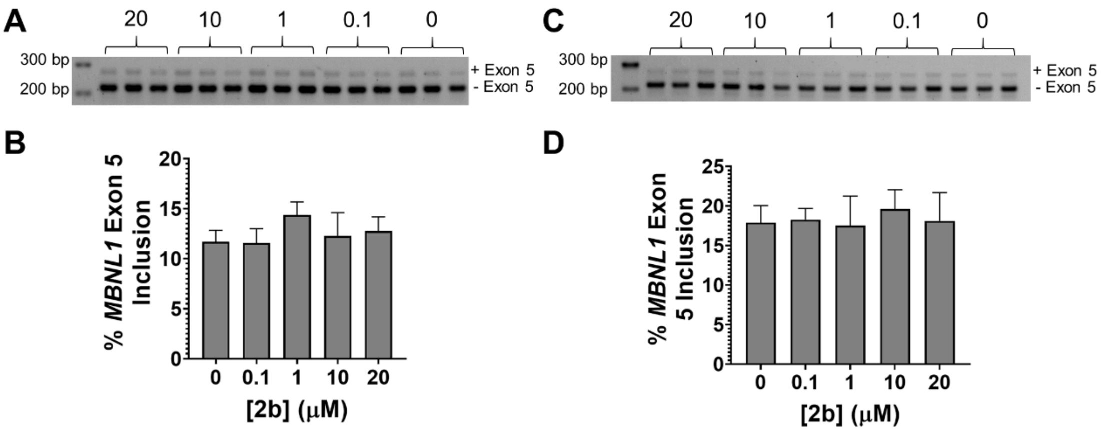


**Figure S6.** Compound **2b** has no effect on *MBNL1* exon 5 splicing in wild-type fibroblasts and myotubes. (A) Representative gel image of *MBNL1* exon 5 splicing in wild-type fibroblasts. (B) Quantification of *MBNL1* exon 5 splicing in wild-type fibroblasts. (C) Representative gel image of *MBNL1* exon 5 splicing in wild-type myotubes. (D) Quantification of *MBNL1* exon 5 splicing in wild-type myotubes. For both panels, n = 3 and error bars indicate SD.

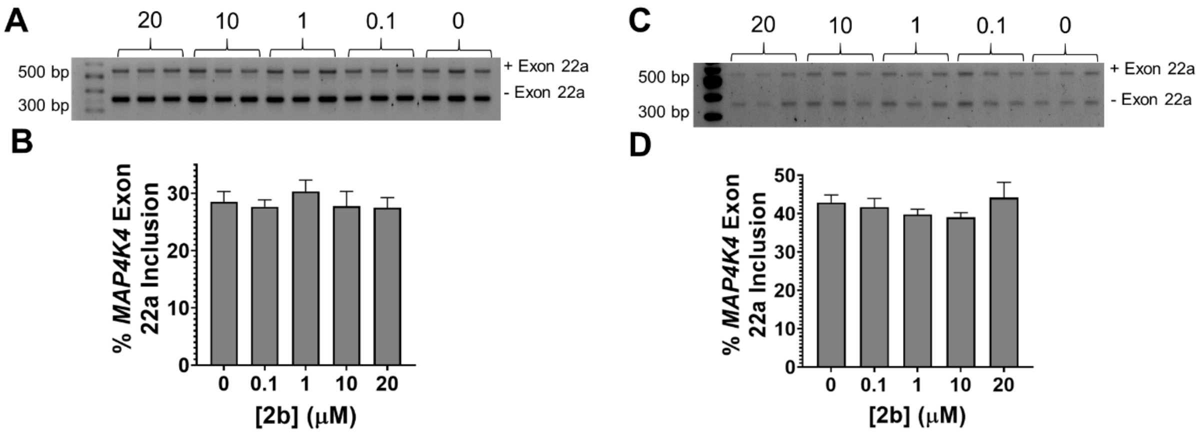


**Figure S7.** Compound **2b** does not affect NOVA-regulated *MAP4K4* splicing in DM1 fibroblasts and myotubes. (A) Representative gel image of *MAP4K4* exon 22a splicing in DM1 fibroblasts treated with **2b**. (B) Quantification of *MAP4K4* exon 22a splicing in DM1 fibroblasts. (C) Representative gel image of *MAP4K4* exon 22a splicing in DM1 myotubes treated with **2b**. (D) Quantification of *MAP4K4* exon 22a splicing in DM1 myotubes. For both panels, n = 3 and error bars indicate SD.

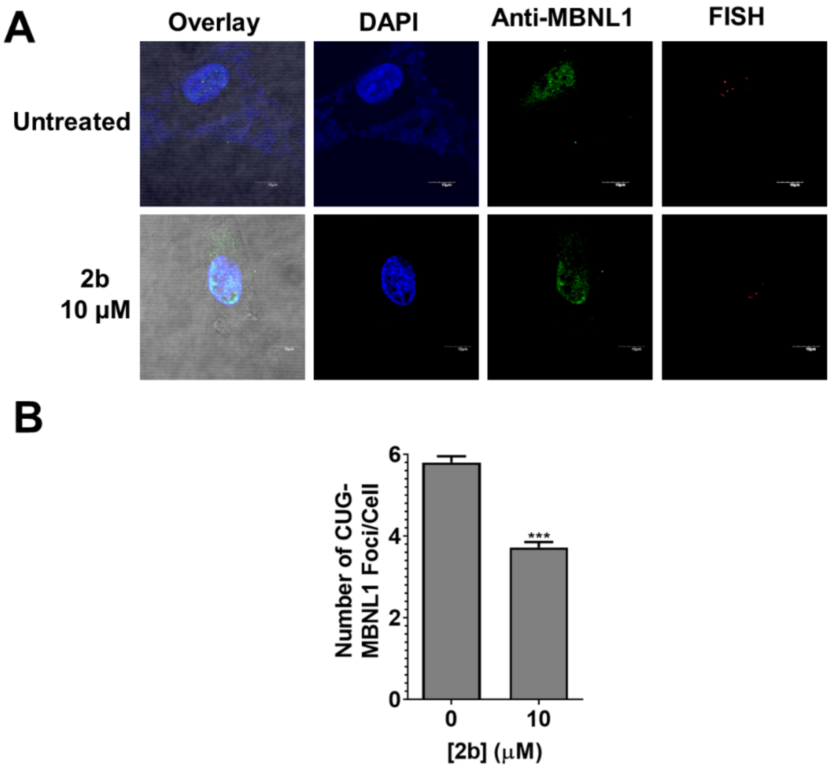


**Figure S8.** Compound **2b** reduces the number of r(CUG)$^{exp}$-containing nuclear foci in DM1 fibroblasts. (A) Representative images of r(CUG)$^{exp}$-MBNL1 foci as determined by MBNL1 immunostaining and RNA FISH. (B) Quantification of the number of nuclear foci/cell (n = 3, 40 cells counted/replicate). Error bars indicate SD; ***, $P < 0.001$, as determined by a two-tailed Student *t*-test.

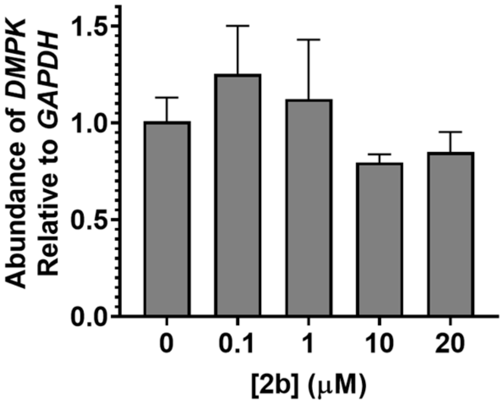


**Figure S9.** Compound **2b** does not affect *DMPK* abundance in DM1 fibroblasts, as determined by RT-qPCR (n = 3). Error bars indicate SD.

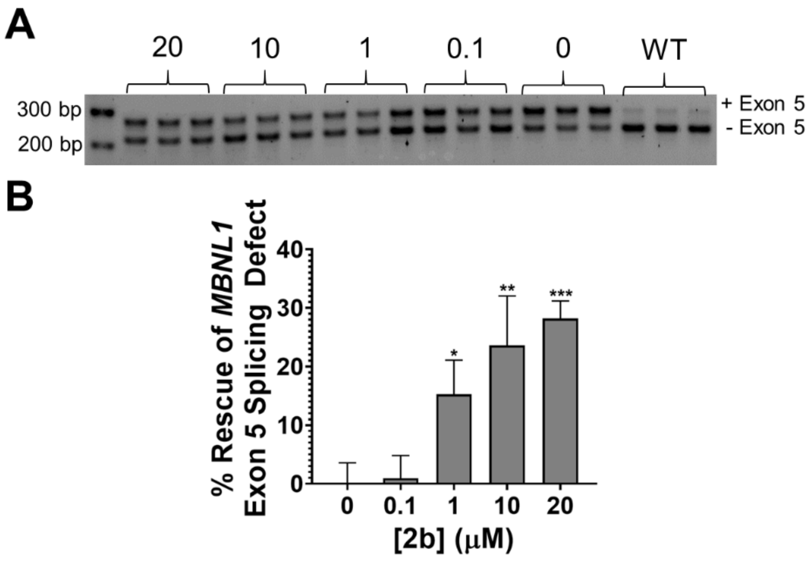


**Figure S10.** Compound **2b** rescues *MBNL1* exon 5 splicing dose-dependently in DM1 patient-derived myotubes. (A) Representative gel image of *MBNL1* exon 5 splicing in DM1 myotubes treated with **2b** (n = 3). (B) Quantification of *MBNL1* exon 5 splicing defect rescue in DM1 myotubes treated with **2b**. Error bars represent SD. *, $P < 0.05$; **, $P < 0.01$; ***, $P < 0.001$, as determined by a one-way ANOVA.

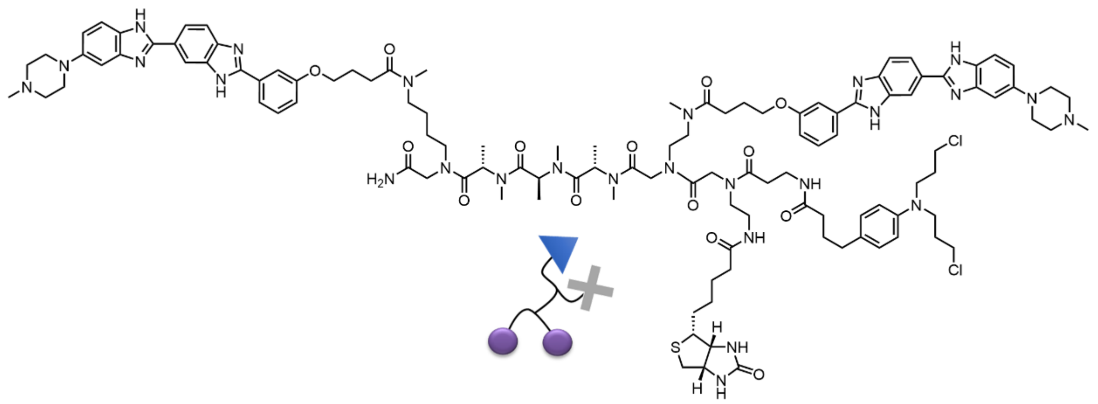

**Figure S11.** Chemical structure of Chem-CLIP probe **2H-K4NMeS-CA-Biotin** synthesized as previously described (1).

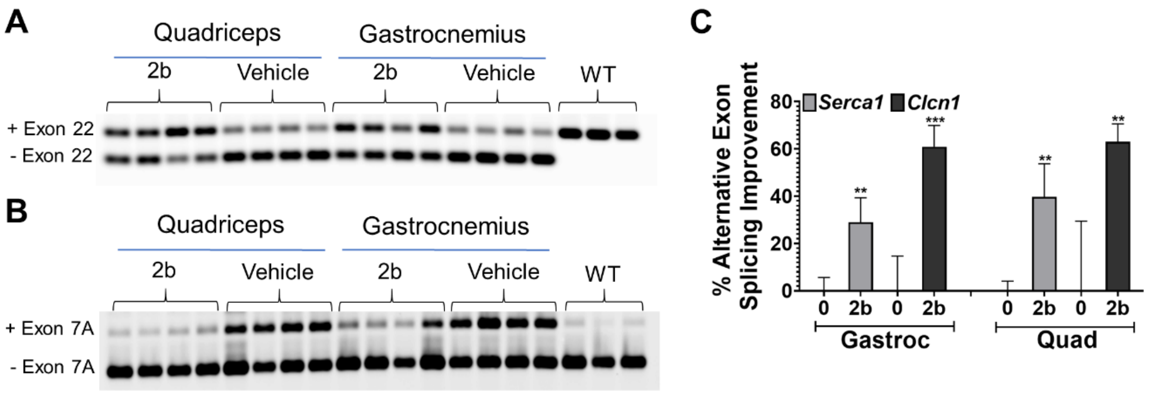


**Figure S12.** Compound **2b** rescues DM1-associated splicing defects in *HSA*$^{LR}$ mice. (A) Representative gel image of *Serca1* exon 22 splicing in the quadriceps and gastrocnemius muscles of *HSA*$^{LR}$ mice treated with **2b** (40 mg/kg). (B) Representative gel image of *Clcn1* exon 7A splicing in the quadriceps and gastrocnemius muscles of *HSA*$^{LR}$ mice treated with **2b**. (B) Quantification of *Serca1* exon 22 and *Clcn1* exon 7A splicing improvement (n = 4 mice per treatment group). Error bars represent SD. **, $P < 0.01$; ***, $P < 0.001$; as determined by a two-tailed Student t-test relative to vehicle treated (0).

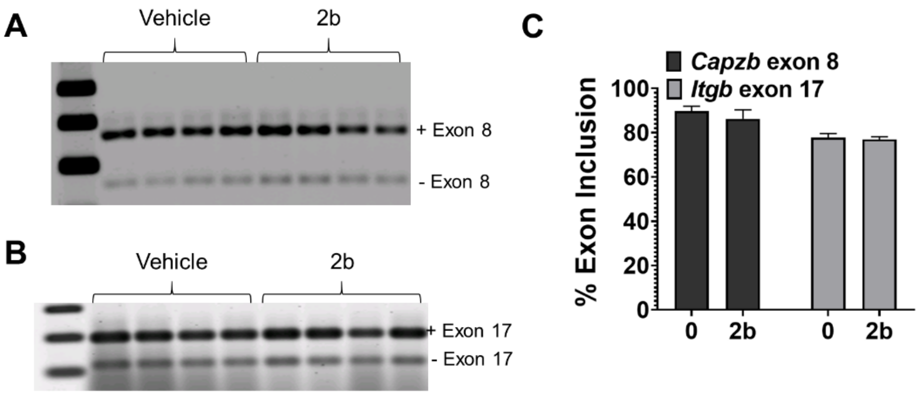


**Figure S13.** Compound **2b** does not affect non-MBNL1 regulated splicing events in *HSA*LR mice (2). (A) Representative gel image of *Capzb* exon 8 splicing in the quadriceps muscle of *HSA*LR mice treated with **2b** (40 mg/kg). (B) Representative gel image of *Itgb* exon 17 splicing in the quadriceps muscle of *HSA*LR mice treated with **2b**. (B) Quantification of *Capzb* and *Itgb* exon inclusion (n = 4 mice/group). Error bars represent SD.

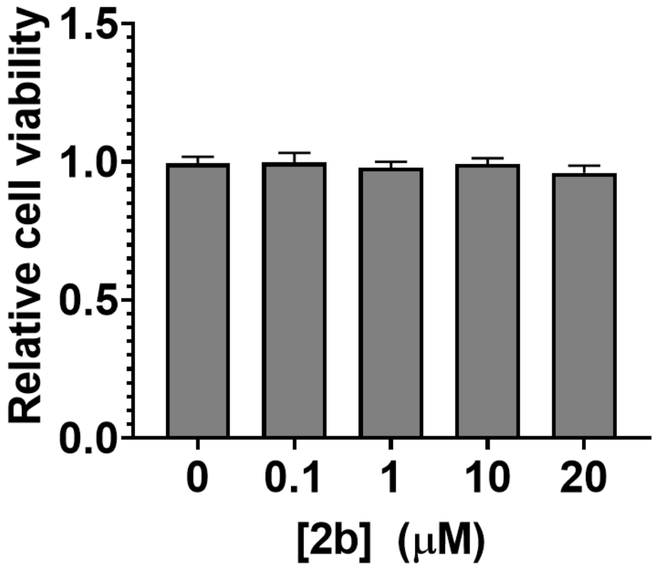


**Figure S14.** Toxicity of **2b** in FECD cells (F35T), as determined by CellTiter-Glo (n = 5). Error bars represent SD.

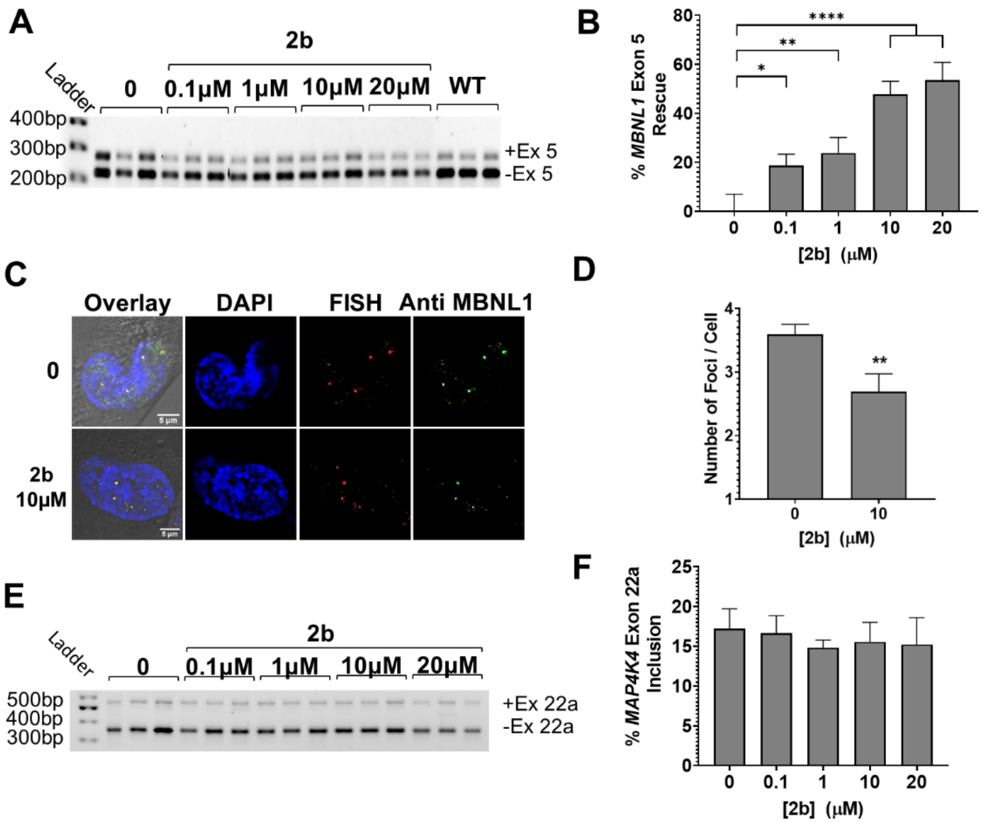


**Figure S15.** Compound **2b** alleviates FECD-associated defects in patient-derived cells. (A) Representative gel image of *MBNL1* exon 5 splicing in FECD (F35T) relative to WT (F20T). (B) Quantification of MBNL1 exon 5 splicing (n =3); *, $P < 0.05$; **, $P < 0.01$; ****, $P < 0.0001$, as determined by a one-way ANOVA. (C) Representative images of r(CUG)$^{exp}$-MBNL1 foci as determined by MBNL1 immunostaining and RNA FISH. (D) Quantification of the number of nuclear foci/cell (n = 3, 40 nuclei counted/replicate); **, $P < 0.01$, as determined by a two-tailed Student *t*-test. (E) Representative gel image of *MAP4K4* exon 22a (non-MBNL1 regulated) splicing in FECD cells treated with **2b**. (F) Quantification of *MAP4K4* exon 22a splicing (n =3). Error bars indicate SD for all plots.

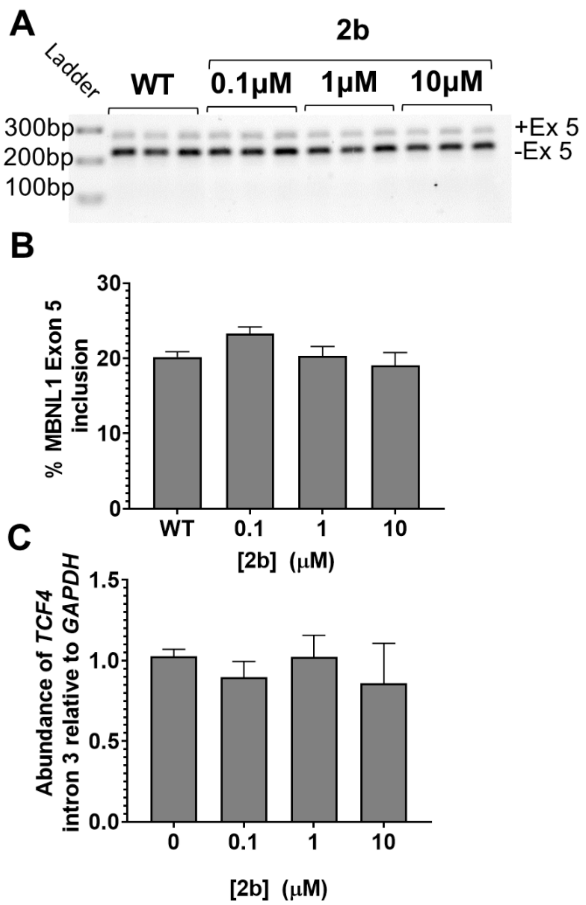


**Figure S16.** Compound **2b** does not affect splicing or *TCF4* intron 3 levels in wild-type corneal endothelium cells without repeat expansion (F20T). (A) Representative gel image of *MBNL1* exon 5 splicing in F20T cells. (B) Quantification of *MBNL1* exon 5 splicing. (C) Abundance of *TCF4* intron 3 levels in F20T cells treated with **2b** (n = 3). Error bars represent SD for both panels B and C.

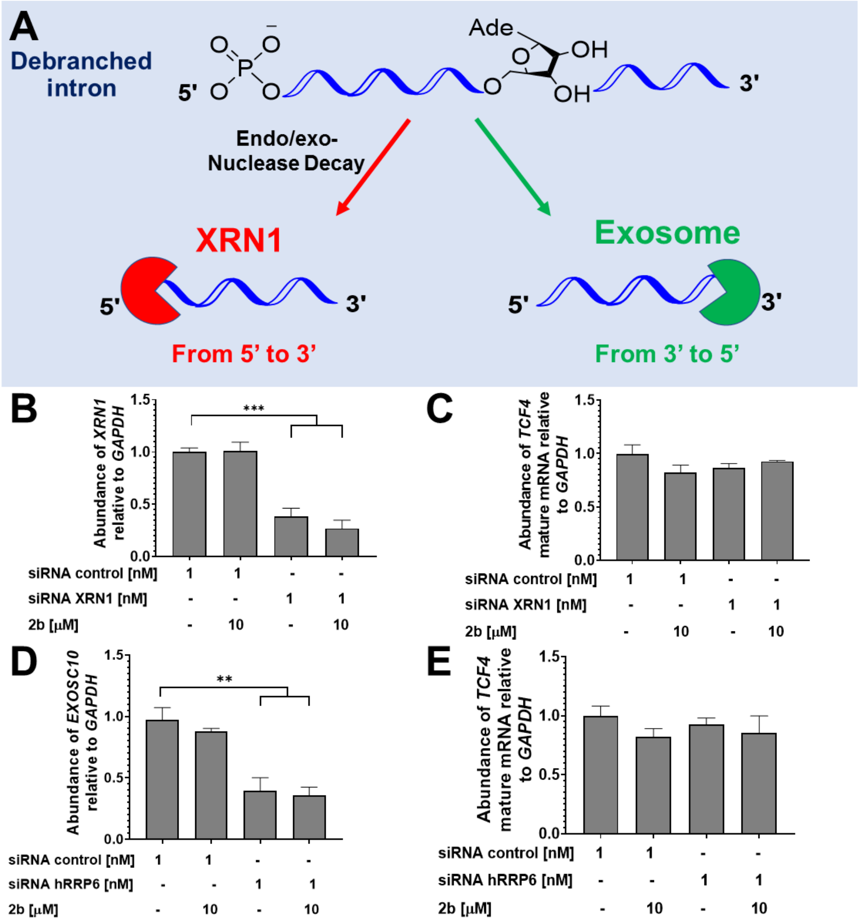


**Figure S17.** Studying the RNA quality control pathways responsible for intron degradation upon **2b** treatment in FECD cells. (A) Decay mechanisms for the Exosome and XRN1 complexes. (B, C) Impact of the siRNA knock down of *XRN1* on **2b**'s ability to reduce *TCF4* intron 3 levels in FECD cells. (B) *XRN1* mRNA levels, as determined by RT-qPCR. (C) *TCF4* mature mRNA levels, as determined by RT-qPCR. (D, E) Impact of the siRNA knock down of *hRRP6* on **2b**'s ability to reduce *TCF4* intron 3 levels in FECD cells. (D) *EXOSC10* (*hRRP6*) mRNA levels, as determined by RT-qPCR. (E) *TCF4* mature mRNA levels, as determined by RT-qPCR. For all panels, n = 3 and error bars represent SD. **, $P < 0.01$; ***, $P < 0.001$, as determined by one-way ANOVA compared to untreated cells ("0").

**Methods:**

**TR-FRET assay to measure r(CUG)$^{exp}$-MBNL1 complex inhibition *in vitro*:** *In vitro* activity of compounds was assessed by inhibition of r(CUG)$^{exp}$-MBNL1 complex formation using a previously reported TR-FRET assay (3, 4). Briefly, biotinylated r(CUG)$_{12}$ (final concentration 80 nM) was folded in 1× Folding Buffer (20 mM HEPES, pH 7.5, 100 mM KCl, and 10 mM NaCl) at 60 °C for 5 min and slowly cooled to room temperature. The buffer was then adjusted to 1× TR-FRET Buffer (1× Folding Buffer supplemented with 2 mM $MgCl_2$, 2 mM $CaCl_2$, 5 mM DTT, 0.1% BSA, 0.05% Tween-20), and the small molecules were added. The solutions were incubated for 1 h at room temperature followed by addition of MBNL1-His$_6$ (60 nM final concentration). The samples were incubated for 15 min, after which 1 µL of antibody solution (1:1 mixture of 8.8 ng/µl Anti-His$_6$-Tb and 800 nM streptavidin XL-665) was added such that the final volume of each sample was 10 µL. The samples were incubated for an additional 15 min at room temperature, followed by measuring the fluorescence of Tb and the FRET between TB and XXL-665. The ratio of fluorescence intensity of 545 and 665 nm as compared to the ratios in the absence of small molecule and in the absence of RNA were used to calculate percent inhibition. The resulting curves were fit to equation 1 to determine $IC_{50}$ values:

$$y = B + \frac{A-B}{1+\left(\frac{IC50}{x}\right)^{hillslope}} \qquad \text{Eq. 1}$$

where y is ratio of fluorescence intensities at 545 nm and 665 nm (F545/F665), x is the concentration of small molecule, B is the maximum observable FRET signal (F545/F665; sample containing RNA and protein but no small molecule added); A is the minimum observable FRET signal (F545/F665; sample containing antibodies but no RNA, protein, or small molecule); and the $IC_{50}$ is the concentration of small molecule where half of the protein is inhibited from binding r(CUG)$_{12}$ by small molecule.

**Microscale thermophoresis (MST) (5):** MST measurements were performed on a Monolith NT.115 system (NanoTemperTechnologies) with Cy5-labeled AT-rich DNA hairpin (5'-Cy5-CGCGAATTCGCGTTTTCGCGAATTCGCG; IDT), Cy5-labeled r(CUG)$_{12}$ (5'-Cy5-GCG(CUG)$_{12}$CGC; Dharmacon), Cy5-labeled r($G_4C_2$)$_8$ (Dharmacon), Cy5-labeled r(CAG)$_{12}$ (5'-Cy5-GCG(CAG)$_{12}$CGC; Dharmacon),and Cy5-labeled base pair control (BP) (5'-Cy5-GCG(CUG)$_6$(CAG)$_6$CGC; Dharmacon), which were deprotected according to the manufacturer's protocol. RNA or DNA (5 nM) was prepared in 1× MST Buffer (8 mM $Na_2HPO_4$, 185 mM NaCl, 1mM EDTA) and folded by heating at 60 °C for 5 min and slowly cooling to room temperature. After cooling, Tween-20 was added to a final concentration of 0.05% (v/v). Compound **2b** was then added to a final concentration of 5 μM, followed by 1:1 dilutions in 10 μL of 1× MST Buffer. Nucleic acid and compound were then mixed 1:1 (v/v). Samples were incubated for 30 min at room temperature and then loaded into premium capillaries (NanoTemper Technologies). The following parameters were used: 5 – 20 % LED, 80% MST power, Laser-On time = 30 s, Laser-Off time = 5 s. Fluorescence was detected using excitation wavelengths of 605–645 nm and emission wavelengths of 680–685 nm. The resulting data were analyzed by thermophoresis analysis and fitted by quadratic binding equation in MST analysis software (NanoTemper Technologies). The dissociation constant was then determined using Equation 2. The reported $K_d$ values are an average of three independent set of experiments.

Eq. 2

$$K_d = \frac{unbound + (bound - unbound)}{2} * ([RNA] + [\mathbf{2b}] + K_d\sqrt{([RNA] + [\mathbf{2b}] + K_d)^2 - 4([RNA] * [\mathbf{2b}])}$$

**Computational modeling methods:** To study the binding of **2b** to r(CUG)$^{exp}$, a previously reported methodology was employed (6). Briefly, a model RNA duplex, r(5'-CCGC<u>U</u>GCGG-3')$_2$, was generated using the nucgen module of AMBER 16 (7). Amber99 (8) force field with revised χ (9) and α/γ torsional parameters taken from the parmbsc0 force field (10) was utilized to describe the RNA. Watson-Crick (WC) base pairing, torsional, and chirality restraints were additionally used to maintain the A-form geometry whenever necessary. Generalized Amber Force Field (GAFF) (11) was used to describe **2b**. To calculate **2b**'s resp charges, Gaussian09 (12) was utilized to first optimize and then calculate the molecular electrostatic potential (MEP). Furthermore, potential energy surface (PES) scans were performed on several torsions of **2b** to better describe its structural properties.

The bound states of the **2b**-RNA complex were investigated using the dynamic docking methodology, which was successfully employed in our previous studies (6). The details of the binding methodology is as follows: All simulations were conducted using the Amber 16 MD package under the Generalized Born implicit solvent model (13). First, we placed the compound 40 Å away from the UU internal loop. The distance between the center-of-mass (COM) of the heavy atoms of **2b** and the COM of the heavy atoms of the loop closing base pairs was used as the reaction coordinate for conducting the dynamic binding approach. The compound was gradually moved towards the internal loop along the reaction coordinate by decrements of 1 Å until the distance reached 0 Å. Then, compound was moved away from the loop along the reaction coordinate by increments of 1 Å until the distance was 40 Å. While the compound was moving towards the UU internal loop, WC base pairing, torsional, and chirality restraints were imposed on the RNA residues except the loop residues to maintain the A-form geometry. This restraint set was called 'flex-restraints' that allowed the loop residues to move naturally while interacting with the compound. While the compound was moved away from the loop region, WC base pairing, torsional, and chirality restraints were imposed on all the RNA residues to transform the RNA structure back to its apo form as observed by NMR and X-ray spectroscopy (14-23). We repeated this process 50 times sequentially to produce 50 initial bound states of the **2b**-RNA complex. These initial bound states were then utilized to conduct implicit-solvent MD simulations. A modified 'flex-restraints' were imposed on the system in the MD simulations where an extra restraint along the reaction coordinate was added to ensure **2b** did not leave the binding site. Each MD simulation was run for 120 ns yielding a total of 6 μs combined MD trajectory, which was used in the cluster analysis. An in-house code was used to calculate the root-mean-square deviation (RMSD) throughout the trajectory and to cluster snapshots having RMSD <= 1.0 Å. MM-PBSA approach was then utilized on clusters having more than 100 snapshots to predict the binding free energies (24). Lowest binding free energy structure is displayed in Figures 2b and s3.

**siRNA constructs:** All siRNA's constructs were purchased from Dharmacon. The siRNA control is an ON-TARGETplus Non-targeting siRNA #4 (cat: #D-001810-04-05), the siRNA XRN1 is an ON-TARGETplus Human XRN1 #54464 (cat: #J-013754-11-0002), and the siRNA hRRP6 was designed as previously reported (Sense: 5'-G.C.A.A.A.A.U.C.U.G.A.A.A.C.U.U.U.C.C.dT.dT-3' and Antisense: 5'-G.G.A.A.A.G.U.U.U.C.A.G.A.U.U.U.U.G.C.dT.dT-3') (25).

**Cell lines:** Compounds were tested in DM1 patient-derived fibroblasts with 500 CUG repeats (GM03987, Coriell Institute) and wild-type fibroblasts (GM07492, Coriell Institute). Compounds were also tested in two cell lines that can be differentiated into myotubes (26): (i) a DM1 (1300 CUG repeats) conditional MyoD-fibroblast cell line and (ii) a wild-type conditional MyoD-fibroblast cell line (generous gifts from D. Furling; Centre de Recherche en Myologie (UPMC/Inserm/CNRS), Institut de Myologie, Paris, France). For FECD studies, compounds were tested in cell lines that were telomerase transformed from Fuchs Endothelial Corneal Dystrophy (FECD) endothelial cells obtained from patients at the time of corneal transplant surgery (under Johns Hopkins University IRB approved protocol (NA_00023119). The F35T (disease model) cell line has a mutant allele containing ~4500 r(CUG) repeats and a wild-type allele with 21 r(CUG) repeats in length. The F20T (WT) cell line has two alleles measuring the same length of 15 r(CUG) repeats. Triplet repeats in this region of *TCF4* greater than 50 (i.e. 150 bp long) are considered pathogenic for FECD.

**Cell culture and compound treatment:** DM1 fibroblasts and wild-type fibroblasts were maintained in growth medium [1× MEM (Corning), 10% FBS (Sigma), 1% MEM non-essential amino acids (Corning), 1% antibiotic/antimycotic solution (Corning), and 1% Glutagro (Corning)], and cells were treated with compound dissolved in growth medium. Conditional MyoD-fibroblast cell lines(26) were grown in growth medium [1× DMEM (Corning), 10% FBS, 1% antibiotic/antimycotic solution, and 1% glutagro]. MyoD-fibroblasts (~80% confluency) were treated with compound and differentiated in differentiation medium [1× DMEM, 1% antibiotic/antimycotic solution, 0.1 mg/mL iron-transferrin, 0.01 mg/mL insulin (Life Technologies), and 2 μg/mL doxycycline (Sigma)].

For FECD studies, F35T and F20T cells were grown in growth medium [1× OptiMEM-1 (Gibco), 5 ng/ml EGF (Human EGF Cat# J64012), 20 ng/ml NGF (Invitrogen Cat# 13257-019), 20 µg/ml ascorbic acid, 200 mg/l calcium chloride, 0.08% chondroitin sulfate, 50 µg/ml gentamicin, 1% antibiotic/antimycotic solution (Corning), and 8% FBS]. Cells were treated with compound dissolved in growth medium.

**Toxicity analysis:** DM1 fibroblasts were plated in a 96-well plate and treated with compound as described above. After 48 h, medium was removed, and cells were washed once with 1× PBS followed by the addition of 50 µL of growth medium containing 10% WST-1 reagent (Roche). Cells were incubated for 30 min at 37 °C, and then absorbance was measured at 450 nm (signal) and 690 nm (background) using a Molecular Devices SpectraMax M5 plate reader. The corrected absorbance was used to determine cell viability relative to untreated cells.

FECD (F35T) were plated in a 96-well plate and treated with **2b** as described above. After 24 h, cell viability was measured using CellTiter-Glo 2.0 (G9242, Promega Corp) per the manufacturer's protocol. Briefly, 100 µL of CellTiter-Glo was added to each well, the plate was then incubated for 10 min at room temperature, and luminescence was measured using a Molecular Devices SpectraMax M5 plate reader.

**Analysis of pre-mRNA splicing:** Rescue of disease-associated splicing defects by a small molecule was completed as previously described (1, 27). Briefly, cells were plated in 12-well plates and treated with compound for 48 h as described above. After 48 h, the cells were lysed, and total RNA was harvested using a Zymo Quick RNA Miniprep Kit per the manufacturer's recommended protocol. Approximately 200 ng of total RNA was reverse transcribed with 100 units of SuperScript III reverse transcriptase (Life

Technologies) at 50 °C or qScript cDNA synthesis kit (20 mL total reaction volume, Quanta BioSciences) per the manufacturers' recommended protocols. Next, 2 µL of the RT reaction was subjected to PCR using GoTaq DNA polymerase (Promega). RT-PCR products were observed after 30 cycles of 95 °C for 30 s, 58 °C for 30 s, 72 °C for 1 min and a final extension at 72 °C for 5 min. Products were separated on a 2% agarose gel run at 100 V for 1 h in 1× TBE buffer, visualized by staining with ethidium bromide, and imaged using a Typhoon 9410 variable mode imager. Gels were quantified using ImageJ. Percent rescue of splicing defects was calculated using equation 3.

$$\% \, Rescue = \frac{\% \, exon \, inclusion \, DM1 - \% \, exon \, inclusion \, treated}{\% \, exon \, inclusion \, DM1 - \% \, exon \, inclusion \, WT} * 100 \qquad \text{(Eq. 3)}$$

**Evaluation of r(CUG)$^{exp}$-MBNL1 foci:** RNA-FISH to image nuclear foci was completed as previously described (1). Briefly, cells were grown in a MatTek Poly-D-Lysine coated 96-well glass bottom plate and treated as described above. After 48 h, cells were fixed followed by FISH as previously described using 1 ng/µL DY547-2'OMe-$(CAG)_6$. Immunostaining of MBNL1 was completed as previously described using anti-MBNL1 (EMD Millipore, #MABE70; diluted 1:50), and goat anti-mouse IgG-DyLight 488 conjugate (1:200 dilution). Nuclei were stained using a 1 µg/µL solution of DAPI in 1× DPBS. Cells were imaged in 1× DPBS using an Olympus FluoView 1000 confocal microscope at 100× magnification. The number of r(CUG)$^{exp}$-MBNL1 foci were counted in 40 nuclei/replicate (120 total nuclei counted over three independent samples).

**RT-qPCR analysis:** Levels of *DMPK*, *TCF4*, *EXOSC10*, and *XRN1* mRNAs were measured by RT-qPCR as follows. Cells were plated in 12-well plates and treated as described above. After 48 h, total RNA was extracted, and reverse transcription was performed as described in "**Analysis of pre-mRNA splicing**". A 2 µL aliquot of the RT reaction was used for each primer pair for qPCR (see Table 1) with SYBR Green Master Mix (Life Technologies) performed on a QuantStudio 5, 384-well Block Real-Time PCR System (Applied Biosciences). Relative abundance of each transcript was determined by normalizing to *GAPDH*.

**Target occupancy by using C-Chem-CLIP:** Target binding of small molecules was assessed using DM1 patient-derived fibroblasts. Cells were grown as monolayers in 100 $mm^2$ dishes in growth medium. Once cells were ~80% confluent, they were treated with growth medium containing **2b** or **3b** for 6 h. Then, **2H-K4NMeS-CA-Biotin** (100 nM final concentration) was added to the growth medium. After 48 h, the cells were lysed and total RNA was harvested using Trizol reagent (Life Technologies). Approximately 10 µg of total RNA was incubated with streptavidin-agarose beads (100 µL, >15 µg/mL binding capacity; Sigma) for 1 h at room temperature. Then the beads were washed with 1× PBS, and the bound RNA was eluted by adding 100 µL of 95% formamide containing 10 mM EDTA, pH 8.2 for 20 min at 60 °C. The eluted RNA was cleaned up using a Zymo Quick RNA miniprep kit per the manufacturer's recommend protocol. Approximately 100 ng of RNA was reverse transcribed using a qScript cDNA synthesis kit (10 µL total reaction volume, Quanta BioSciences); 1 µL of the RT reaction was used with each primer set for qPCR with SYBR Green Master Mix (Life Technologies) performed on a 7900HT Fast Real-Time PCR System (Applied Biosystems). Relative abundance was determined by normalizing to *GAPDH*.

**Evaluation of 2b in *HSA*$^{LR}$ mice:** All experimental procedures, mouse handling, and husbandry were completed in accordance with the Association for Assessment and Accreditation of Laboratory Animal Care. A mouse model for DM1, *HSA*$^{LR}$ in line 20b (28), was used. *HSA*$^{LR}$ mice express human skeletal actin RNA with 220 r(CUG) repeats in the 3' UTR. Age- and gender-matched *HSA*$^{LR}$ mice were injected intraperitoneally with either 40 mg/kg of **2b** in water for treatment or the same volume of 0.9% NaCl for vehicle-treated mice once per day for 7 days. Mice were euthanized one day after the last injection, and the quadriceps and gastrocnemius muscle were harvested. RNA was extracted from the tissue, cDNA was synthesized, and PCR was performed as previously described (29).

**Synthetic Methods:**

**General Synthetic Methods:** Microwave reactions were carried out using a Biotage Initiator+ SP Wave microwave. Compounds were purified by preparative reverse phase HPLC using a Waters 1525 Binary HPLC pump equipped with a Waters 2487 dual absorbance detector system and a Waters Sunfire C18 OBD 5 µm 19 x 150 mm column. Absorbance was monitored at 345 and 220 nm. A gradient of 0-100% MeOH in $H_2O$ with 0.1% TFA over 60 min was used for compound purification. Purity was assessed by analytical HPLC using a Waters Symmetry C18 5 µm 4.6 x 150 mm column. Small molecules were analyzed using a gradient of 0-100% methanol in water with 0.1% TFA over 60 min. $^{1}H$ NMR spectra were collected using a Bruker 400 MHz NMR, while $^{13}C$ NMR spectra were collected on a Bruker 600 MHz NMR. High resolution mass spectrometry (HR-MS) was completed on an Applied Biosystems MALDI ToF Analyzer 4800 Plus using an α-cyano-4-hydroxycinnamic acid matrix with calibration standards. All compounds evaluated had ≥95% purity.

**Scheme S-1. Synthesis of 1a and 1b.**

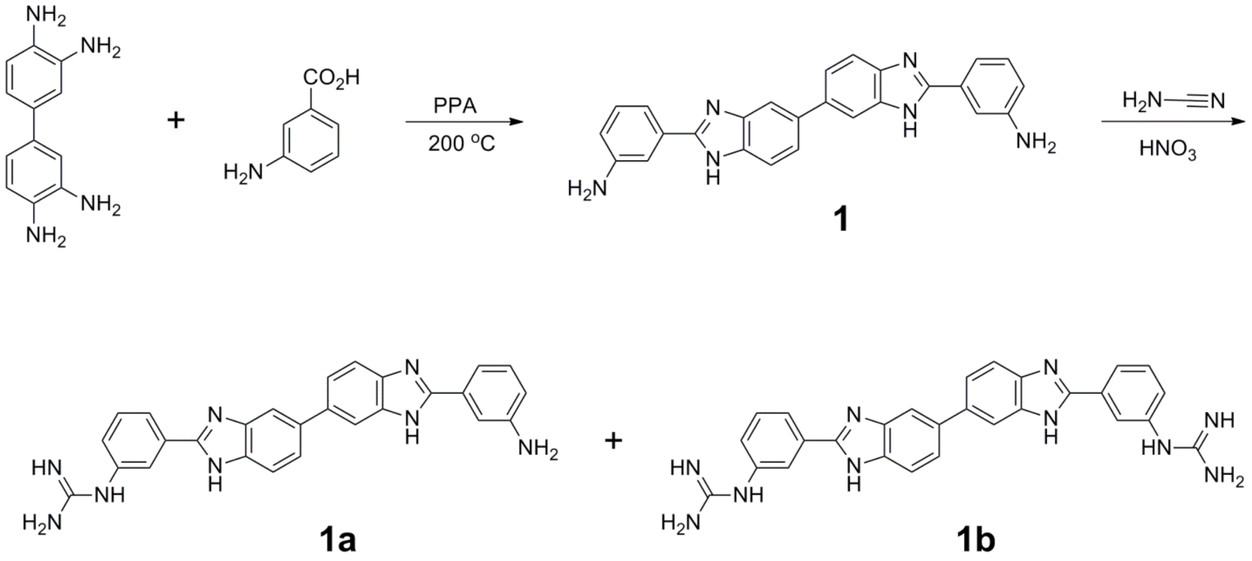

*Synthesis of* ***1a*** *and* ***1b***. Compound **1** (10 mg, 16.4 µmol) (30) was suspended in a mixture of water (1.50 mL) and ethanol (300 µL). Then cyanamide (50 mg, 1.2 mmol) and nitric acid (5 drops) were added, and the reaction mixture was heated to 120 °C via a microwave for 6 h. Then, the reaction mixture was diluted in water and purified by preparative reverse phase HPLC as described in the General Methods, affording 2.56 µmoles of **1a** (16%, light brown solid) and 467 nmoles of **1b** (3%, light brown solid).

**1a:** $^{1}$H NMR (400 MHz, $CD_3OD$): δ 8.10 (m, 2H), 8.02 (d, 1H, $J$ = 1), 7.98 (d, 1H, $J$ = 2), 7.95 (d, 1H, $J$ = 1), 7.93 (d, 1H, $J$ = 2), 7.88 (d, 1H, $J$ = 9), 7.81 (d, 1H, $J$ = 8), 7.72 (m, 2H), 7.50 (m, 1H), 7.41 (m, 3H), 7.07 (m, 1H). HRMS (m/z): $[M]^+$ calcd. for $C_{27}H_{22}N_8$, 459.2040; found, 459.1970.

**1b:** $^{1}$H NMR (400 mHz, $CD_3OD$): δ 8.10 (m, 4H), 7.97 (s, 2H), 7.80 (m, 2H), 7.74 (m, 4H), 7.52 (m, 2H). $^{13}$C NMR (600 mHz, $CD_3OD$): δ 158.63, 138.02, 132.64, 130.13, 127.25, 126.64, 125.28, 118.43, 117.05, 116.52, 116.05, 1114.10. HRMS (m/z): $[M]^+$ calcd. for $C_{28}H_{24}N_{10}$, 501.2258; found, 501.2247.

**Scheme S-2. Synthesis of 2a and 2b.**

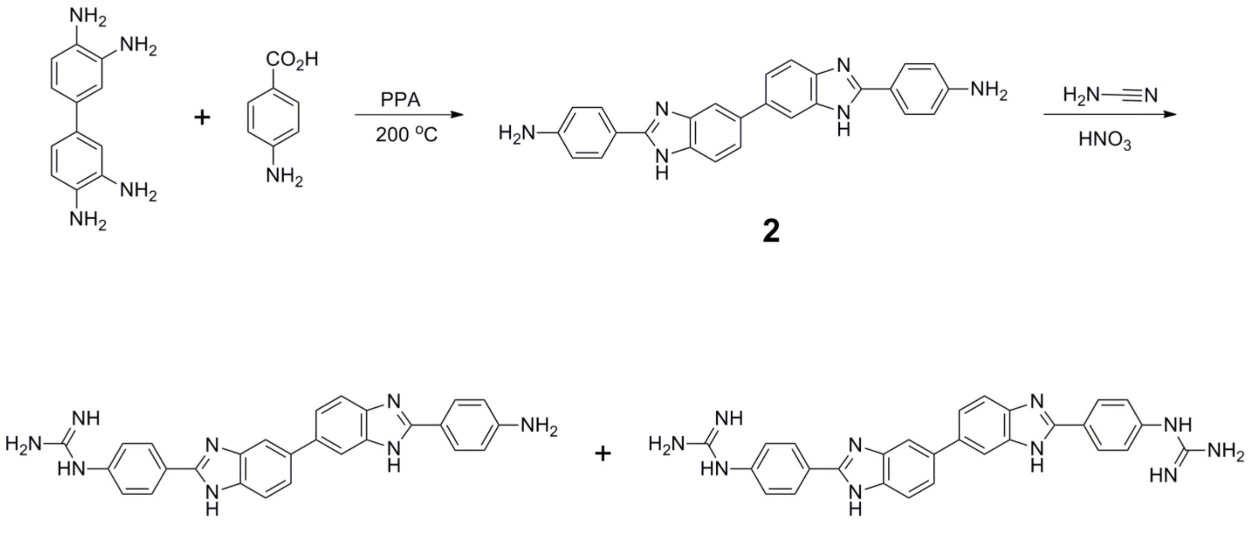


2a 2b

*Synthesis of* ***2a*** *and* ***2b***. Compound **2** (10 mg, 16.4 µmol) (30), was suspended in a mixture of water (1.5 mL) and ethanol (300 µL). Then, cyanamide (50 mg, 1.2 mmol) and nitric acid (5 drops) were added, and the reaction mixture was refluxed overnight. Afterwards, the reaction mixture was diluted in water and purified by preparative reverse phase HPLC as described in the General Methods, affording 1.07 μmoles of **2a** (7%, light brown solid) and 161 nmoles of **2b** (1%, light brown solid).

**2a** $^{1}H$ NMR (400 mHz, $CD_3OD$): δ 8.03-8.01 (m, 2H), 7.94-7.79 (m, 10H), 6.86 (d, 2H, $J$ = 9). HRMS (m/z): $[M]^+$ calcd. for $C_{27}H_{22}N_8$, 459.2040; found, 459.1930.

**2b** $^{1}H$ NMR (400 mHz, $CD_3OD$): δ 8.25 (d, 4H, $J$ = 9), 8.05 (s, 2H), 7.88 (s, 4H) 7.59 (d, 4H, $J$ = 9). $^{13}C$ NMR (600 mHz, $CD_3OD$): δ 157.82, 151.77, 140.80, 139.74, 136.91, 135.77, 130.27, 126.43, 125.99, 124.86, 119.09, 117.72, 115.92, 113.89. HRMS (m/z): $[M]^+$ calcd. for $C_{28}H_{24}N_{10}$, 501.2258; found, 501.2231.

**Scheme S-3. Synthesis of 3b.**

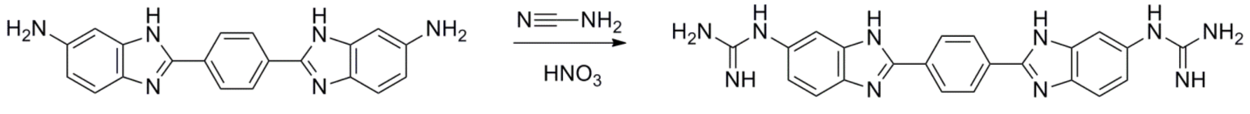


*Synthesis of **3b**.* 2,2′-*p*-Phenylene-bis(5-aminobenzimidazole) (**3**) was synthesized as previously described (31), and 150 mg (0.44 mmol) was suspended in water (1.5 mL)and ethanol (0.5 mL). Cyanamide (730 mg, 17.6 mmol) was added followed by 6 drops of concentrated nitric acid. The reaction was then placed in a Biotage Initiator+ SP Wave microwave at 130°C for 6 h. Next, the reaction mixture was diluted with water and purified by preparative reverse phase HPLC as described in the "**General Methods**" section, affording 50 mg of the desired product (27%, yellow solid). $^{1}H$ NMR (400 mHz, $CD_3OD$): δ 8.36 (s, 4H), 7.79 (d, 2H, $J$ = 9), 7.66 (d, 2H, $J$ = 2), 7.33 (dd, 2H, $J$ = 2, 9). $^{13}C$ NMR (600 mHz, $CD_3OD$): δ 158.61, 152.86, 138.98, 137.17, 132.73, 131.16, 124.09, 117.05, 113.97. HRMS (m/z): $[M]^+$ calcd. for $C_{22}H_{21}N_{10}$, 425.1945; found, 425.1849.

### Compound Characterization:

## 1a:

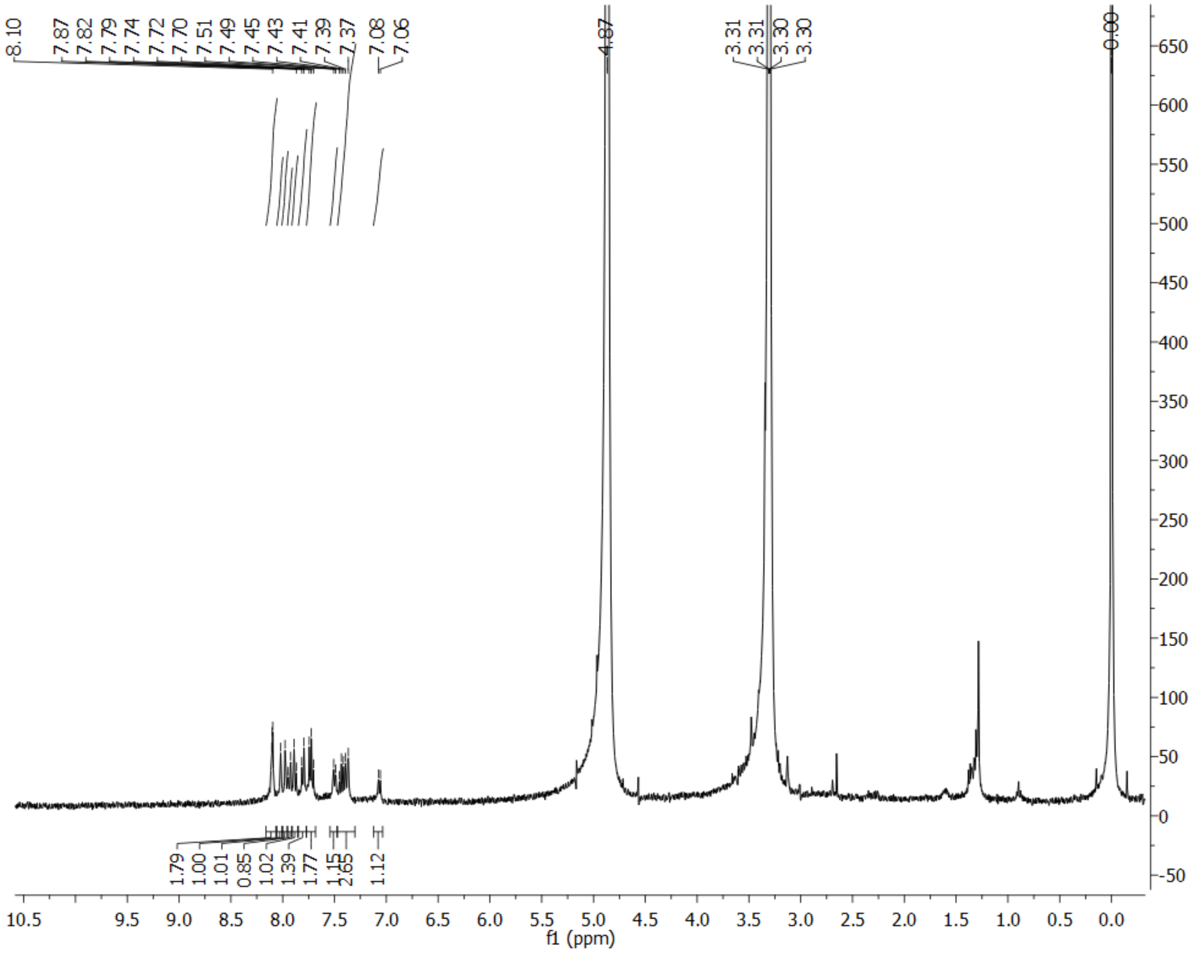

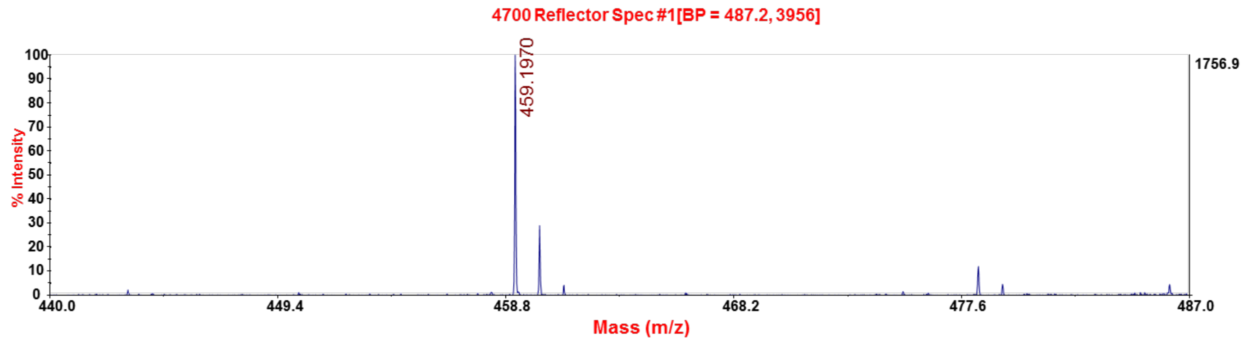

**1b:**

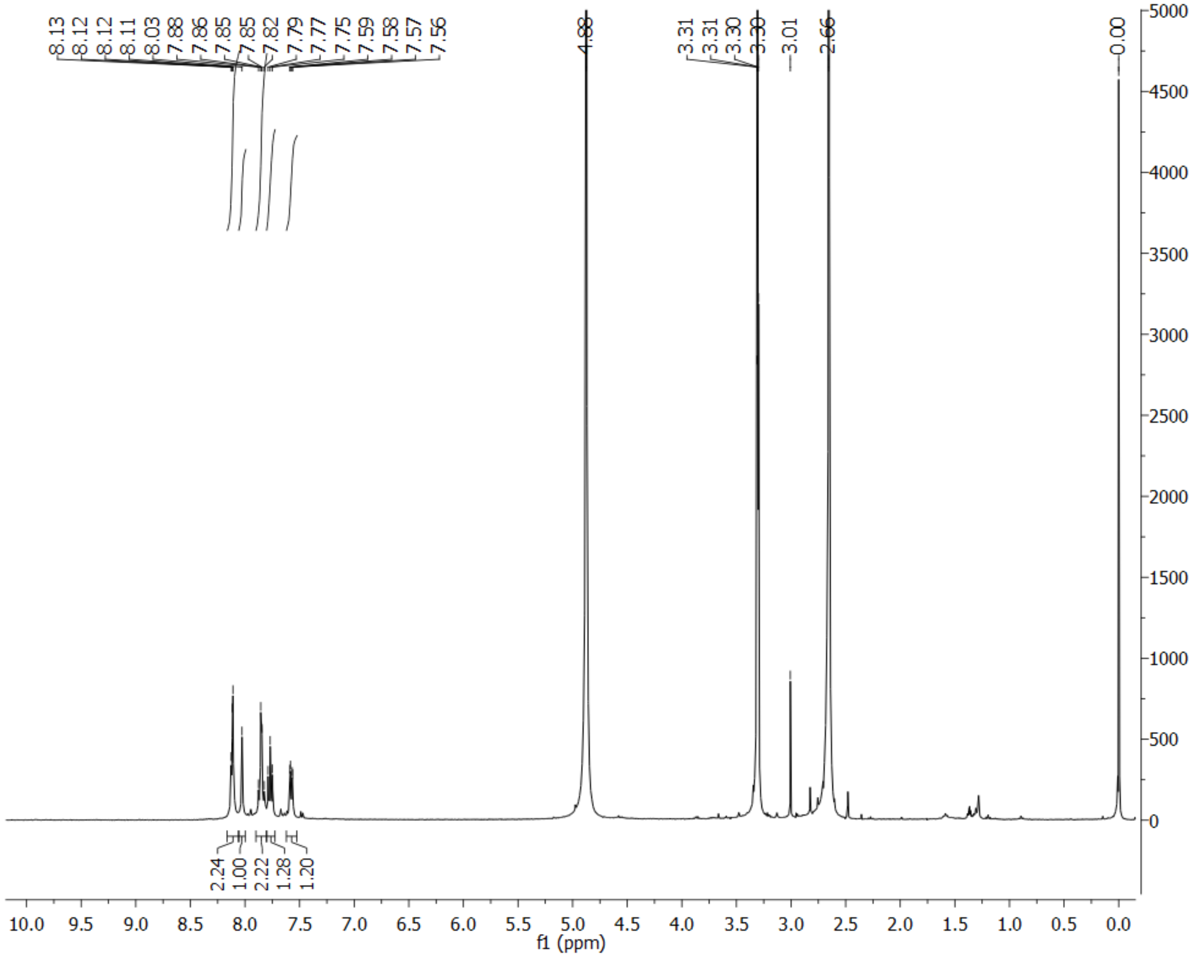

8.13
8.12
8.12
8.11
8.03
7.88
7.86
7.85
7.85
7.82
7.79
7.77
7.75
7.59
7.58
7.57
7.56
4.88
3.31
3.31
3.30
3.30
3.01
2.66
0.00
2.24
1.00
2.22
1.28
1.20
f1 (ppm)


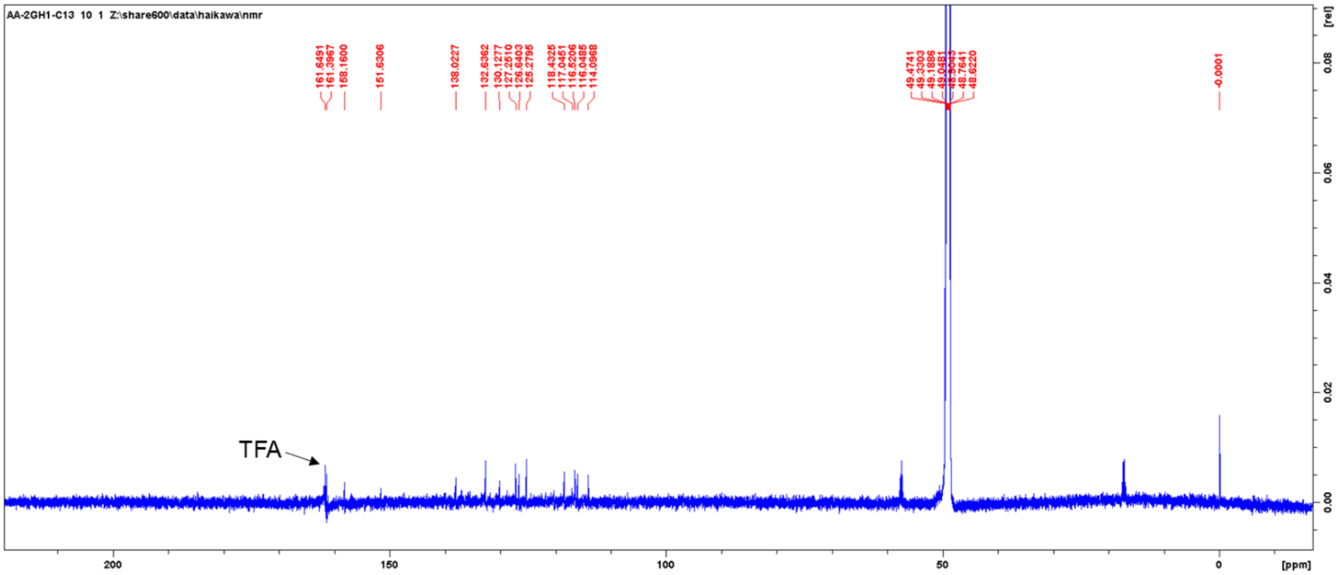

AA-2GH1-C13 10 1 Z:\share600\data\haikawa\nmr
TFA
[ppm]
[rel]

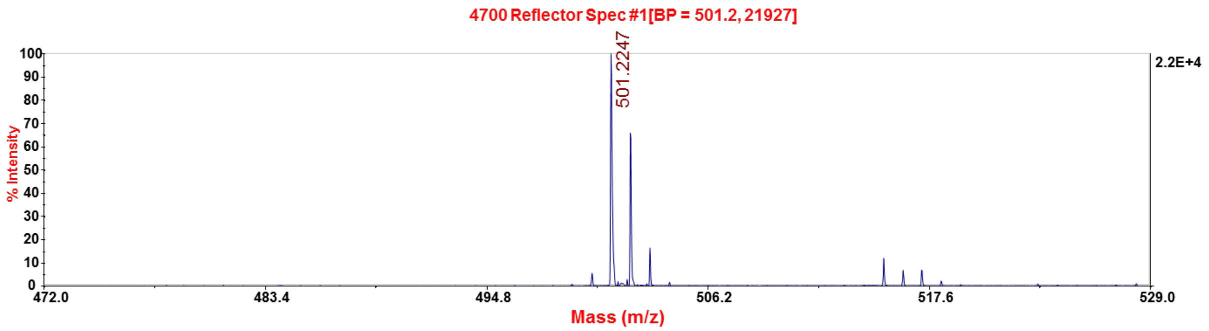
4700 Reflector Spec #1[BP = 501.2, 21927]
501.2247
2.2E+4
% Intensity
Mass (m/z)

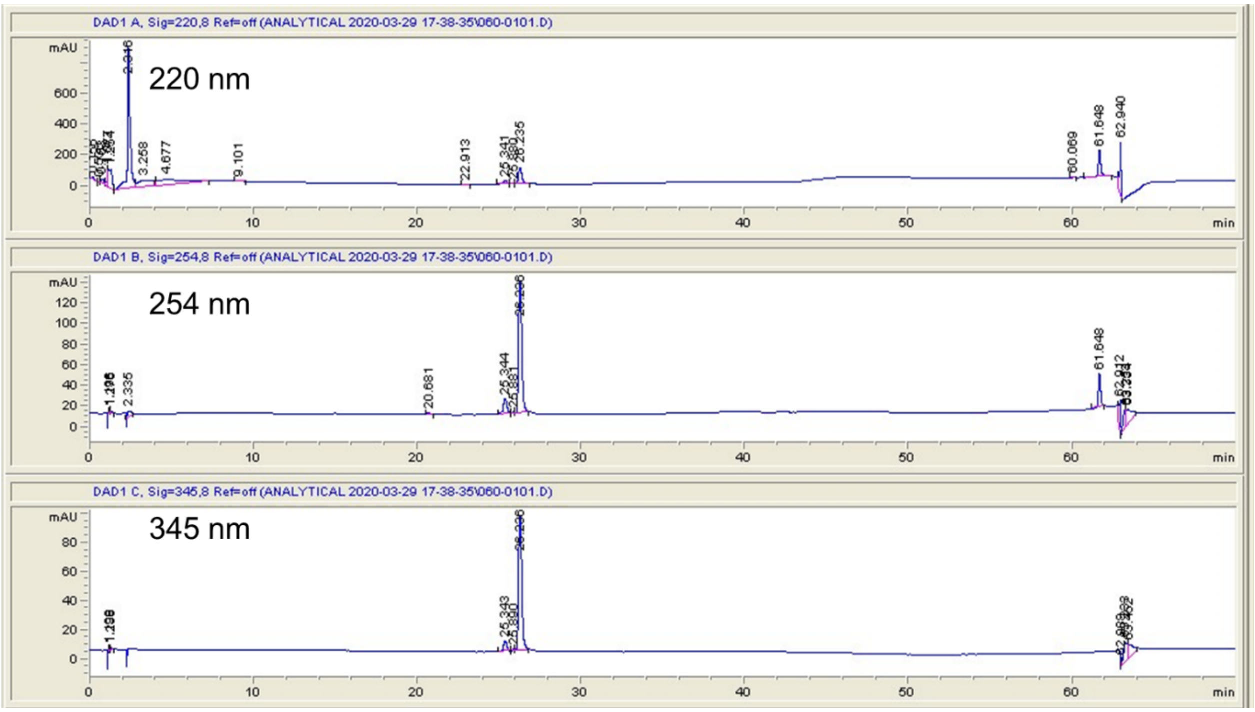
DAD1 A, Sig=220,8 Ref=off (ANALYTICAL 2020-03-29 17-38-35\060-0101.D)
220 nm
DAD1 B, Sig=254,8 Ref=off (ANALYTICAL 2020-03-29 17-38-35\060-0101.D)
254 nm
DAD1 C, Sig=345,8 Ref=off (ANALYTICAL 2020-03-29 17-38-35\060-0101.D)
345 nm
mAU
min

**2a:**

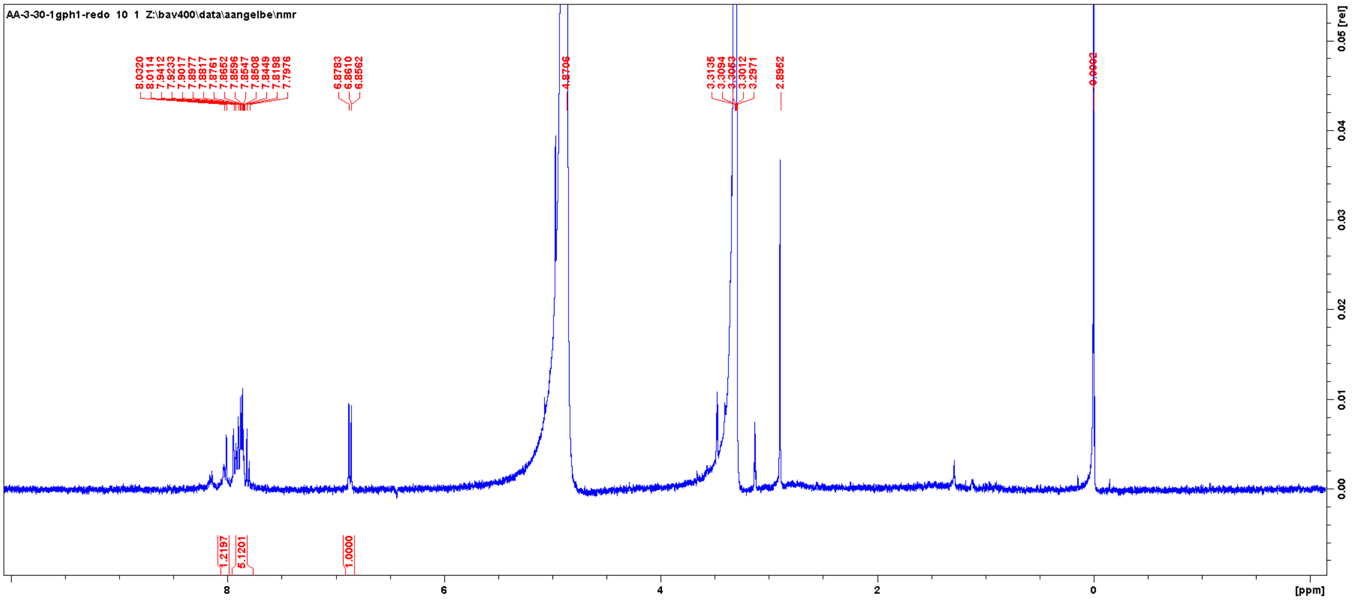
AA-3-30-1gph1-redo 10 1 Z:\bav400\data\aangelbe\nmr
[ppm]
0.05 [rel]

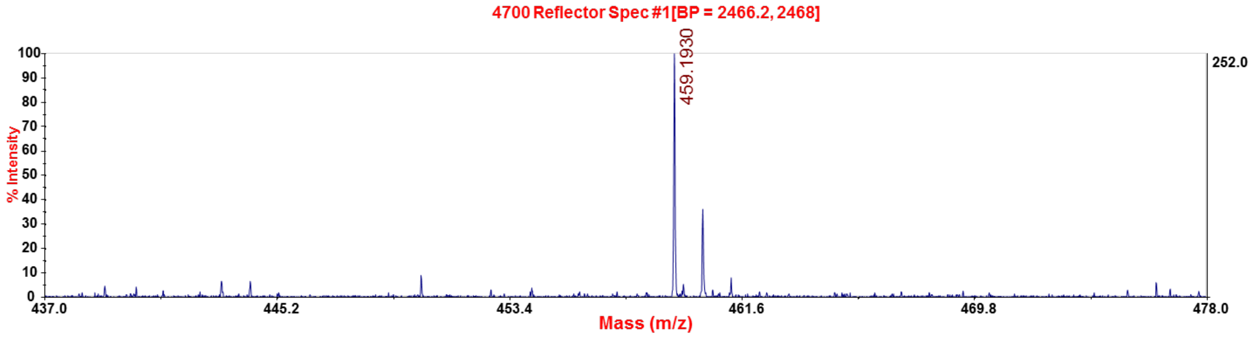
4700 Reflector Spec #1[BP = 2466.2, 2468]
459.1930
% Intensity
Mass (m/z)
252.0

**2b:**

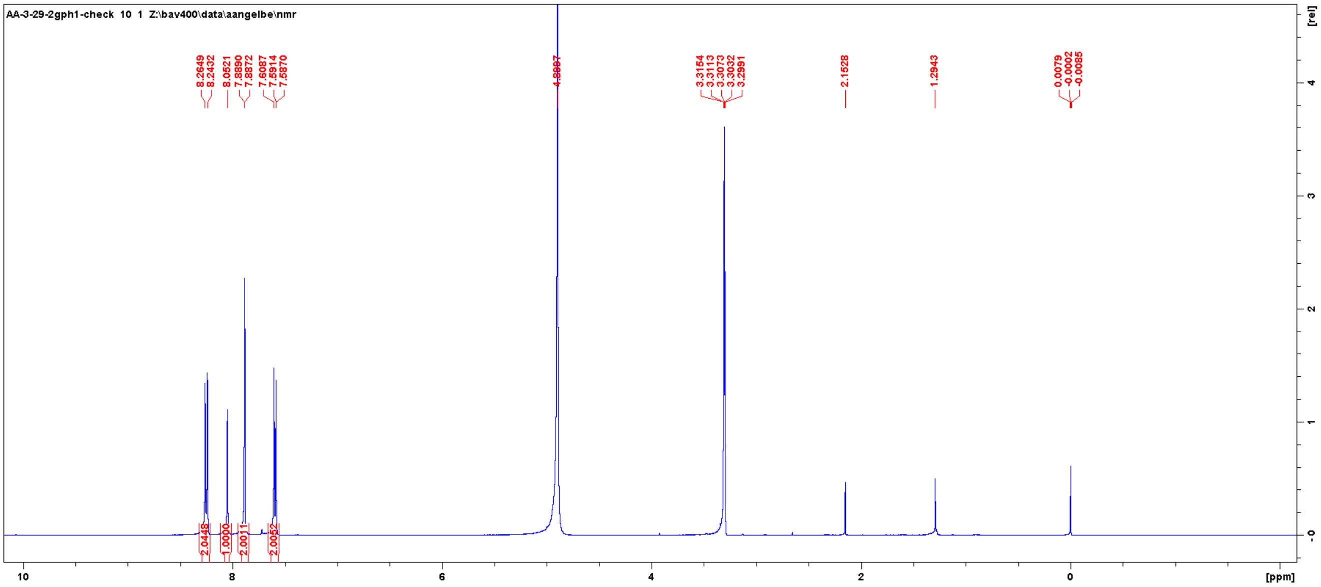

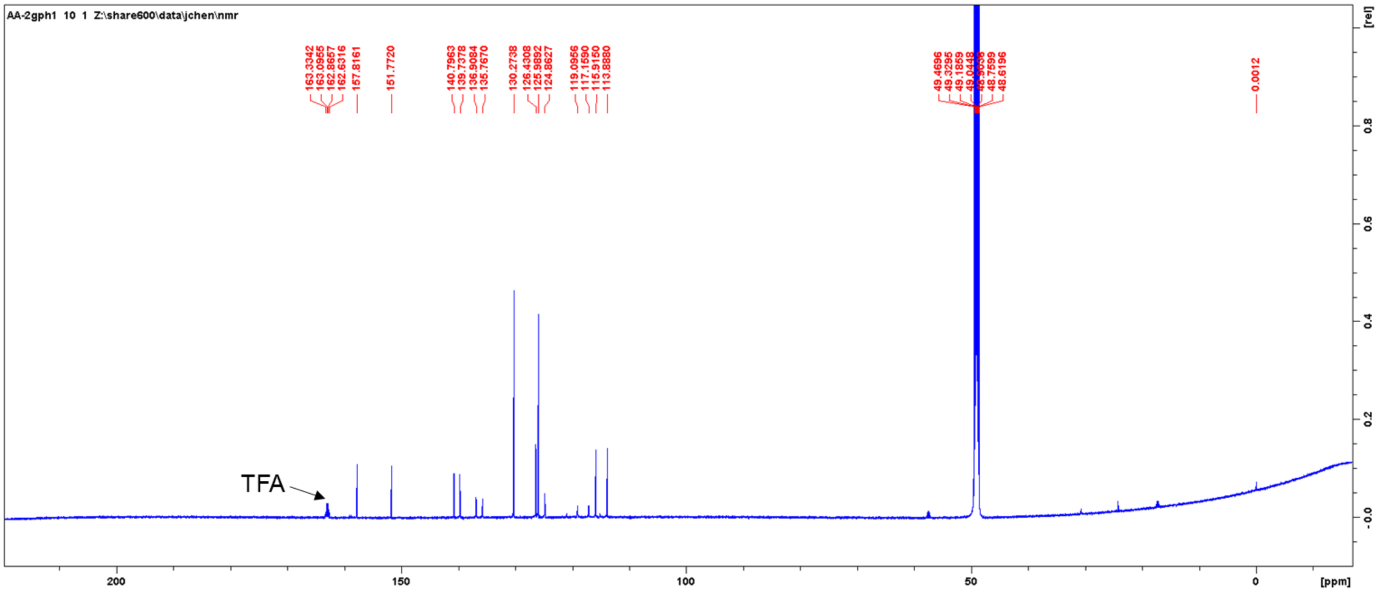
TFA

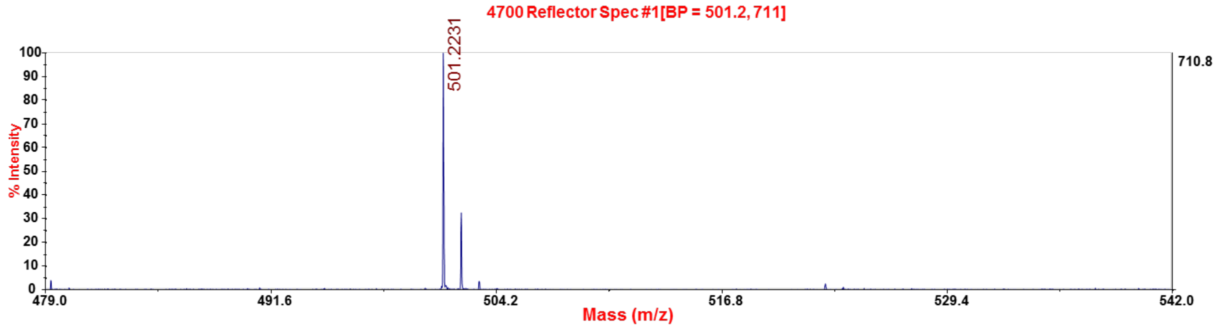
4700 Reflector Spec #1[BP = 501.2, 711]
501.2231
710.8
% Intensity
Mass (m/z)

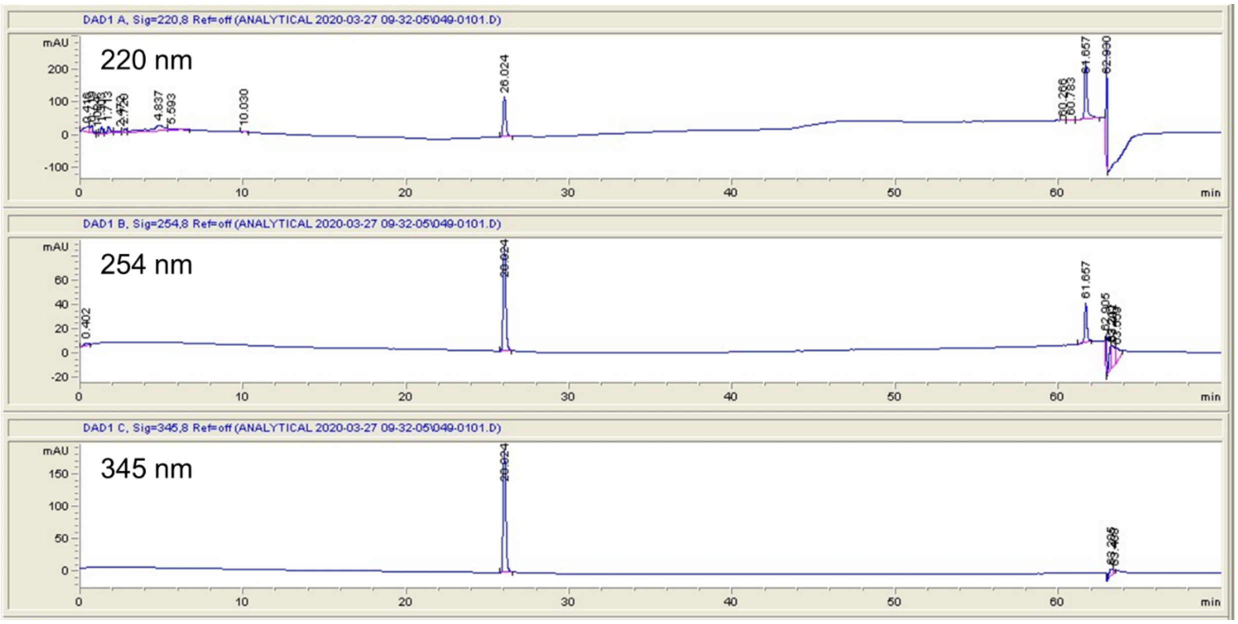

DAD1 A, Sig=220,8 Ref=off (ANALYTICAL 2020-03-27 09-32-05\049-0101.D)
220 nm
26.024
61.657
62.930
DAD1 B, Sig=254,8 Ref=off (ANALYTICAL 2020-03-27 09-32-05\049-0101.D)
254 nm
61.657
DAD1 C, Sig=345,8 Ref=off (ANALYTICAL 2020-03-27 09-32-05\049-0101.D)
345 nm
mAU
min


**3b.**

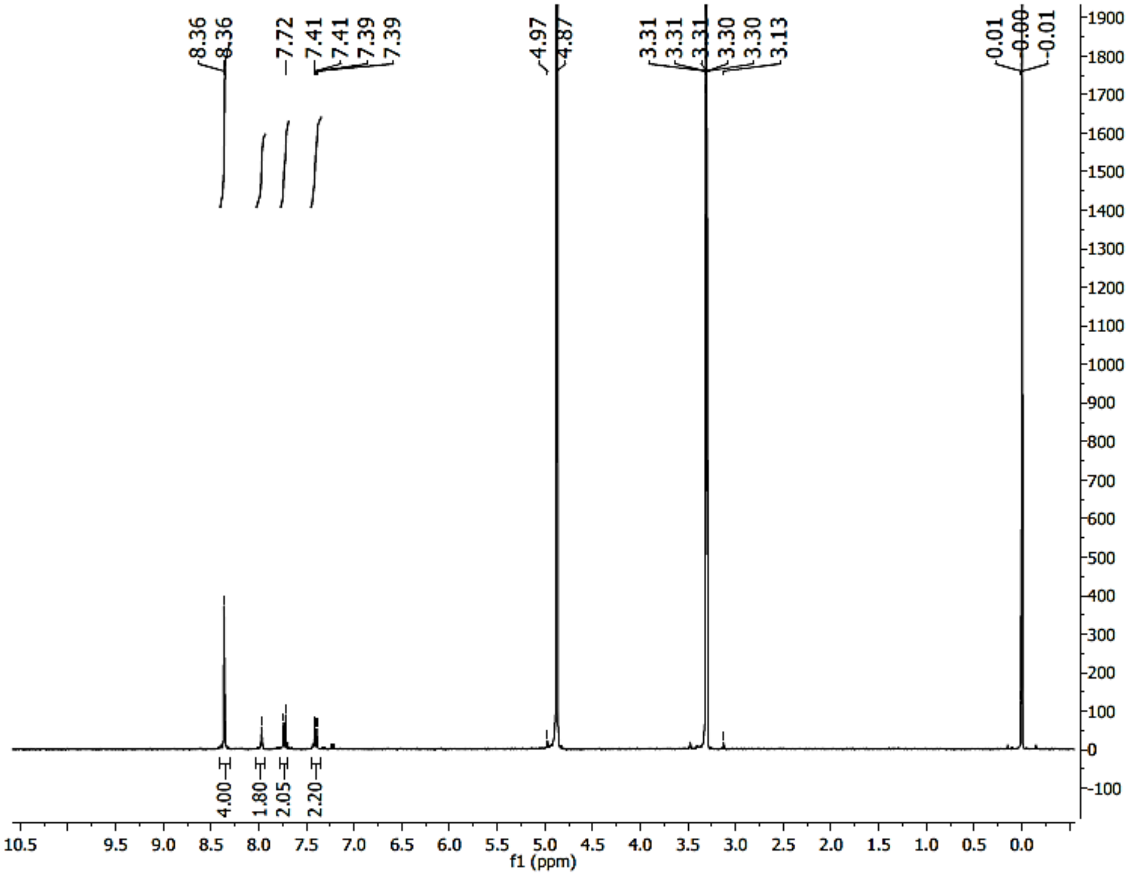

8.36
8.36
7.72
7.41
7.41
7.39
7.39
4.97
4.87
3.31
3.31
3.31
3.30
3.30
3.13
0.01
0.00
-0.01
4.00
1.80
2.05
2.20
f1 (ppm)

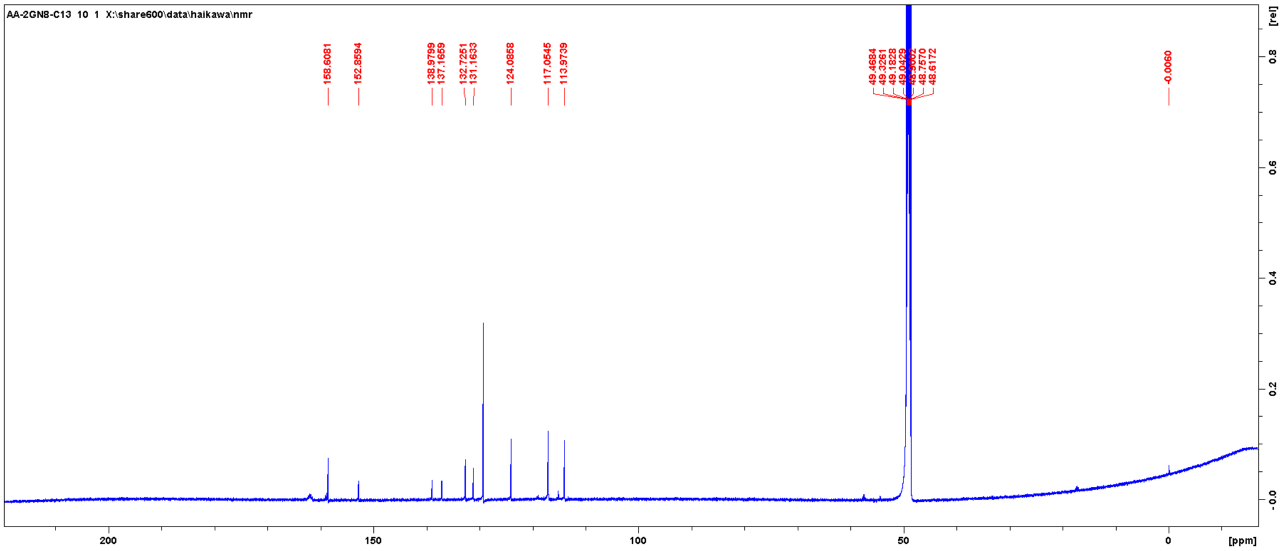
AA-2GN8-C13 10 1 X:\share600\data\haikawa\nmr
158.6081
152.8594
138.9799
137.1659
132.7251
131.1633
124.0858
117.0545
113.9739
-0.0060
[ppm]
[rel]

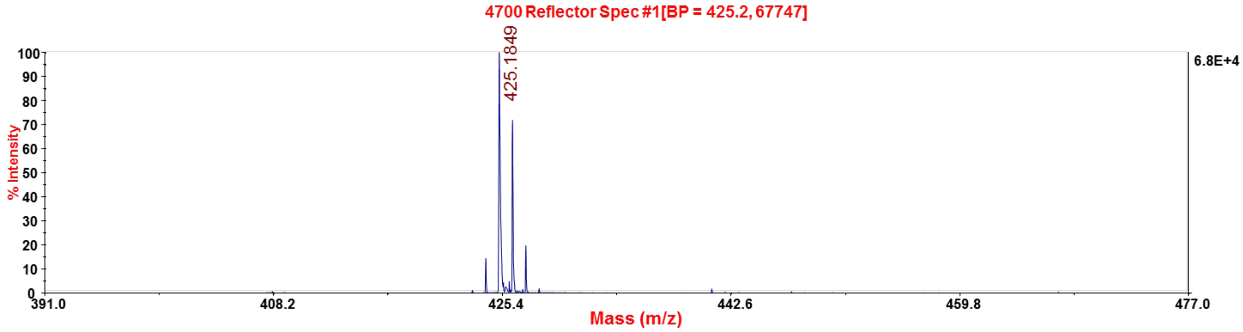
4700 Reflector Spec #1[BP = 425.2, 67747]
425.1849
6.8E+4
% Intensity
Mass (m/z)

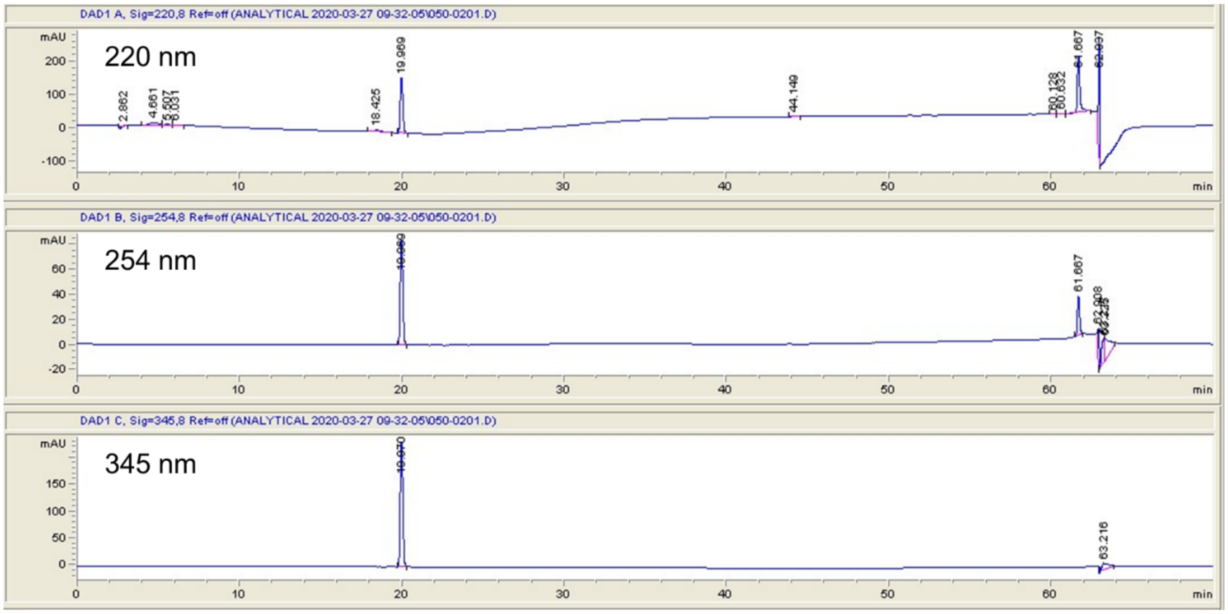
DAD1 A, Sig=220,8 Ref=off (ANALYTICAL 2020-03-27 09-32-05\050-0201.D)
220 nm
DAD1 B, Sig=254,8 Ref=off (ANALYTICAL 2020-03-27 09-32-05\050-0201.D)
254 nm
DAD1 C, Sig=345,8 Ref=off (ANALYTICAL 2020-03-27 09-32-05\050-0201.D)
345 nm
mAU
min

**References:**


1. S. G. Rzuczek *et al.*, Precise small-molecule recognition of a toxic CUG RNA repeat expansion. *Nat. Chem. Biol.* **13**, 188-193 (2017).
2. X. Lin *et al.*, Failure of MBNL1-dependent post-natal splicing transitions in myotonic dystrophy. *Hum. Mol. Genet.* **15**, 2087-2097 (2006).
3. C. Z. Chen *et al.*, Two high-throughput screening assays for aberrant RNA-protein interactions in myotonic dystrophy type 1. *Anal. Bioanal. Chem.* **402**, 1889-1898 (2012).
4. R. Parkesh *et al.*, Design of a bioactive small molecule that targets the myotonic dystrophy type 1 RNA via an RNA motif-ligand database & chemical similarity searching. *J. Am. Chem. Soc.* **134**, 4731-4742 (2012).
5. Y. Li, M. D. Disney, Precise small molecule degradation of a noncoding RNA identifies cellular binding sites and modulates an oncogenic phenotype. *ACS Chem. Biol.* **13**, 3065-3071 (2018).
6. J. L. Childs-Disney *et al.*, Structure of the myotonic dystrophy type 2 RNA and designed small molecules that reduce toxicity. *ACS Chem. Biol.* **9**, 538-550 (2014).
7. D.A. Case *et al.*, AMBER 2016. *University of California, San Francisco* (2016).
8. W. D. Cornell *et al.*, A second generation force field for the simulation of proteins, nucleic acids, and organic molecules. *J. Am. Chem. Soc.* **117**, 5179-5197 (1995).
9. I. Yildirim, H. A. Stern, S. D. Kennedy, J. D. Tubbs, D. H. Turner, Reparameterization of RNA $\chi$ torsion parameters for the AMBER force field and comparison to NMR spectra for cytidine and uridine. *J. Chem. Theory Comp.* **6**, 1520-1531 (2010).
10. A. Pérez *et al.*, Refinement of the AMBER force field for nucleic acids: improving the description of $\alpha/\gamma$ conformers. *Biophys. J.* **92**, 3817-3829 (2007).
11. J. Wang, R. M. Wolf, J. W. Caldwell, P. A. Kollman, D. A. Case, Development and testing of a general amber force field. *J. Comput. Chem.* **25**, 1157-1174 (2004).
12. M. Frisch *et al.*, Gaussian 09; Gaussian, Inc: Wallingford, CT (2009).

13. A. Onufriev, D. Bashford, D. A. Case, Exploring protein native states and large-scale conformational changes with a modified generalized born model. *Proteins* **55**, 383-394 (2004).
14. G. E. Chapman, B. D. Abercrombie, P. D. Cary, E. M. Bradbury, The measurement of small nuclear overhauser effects in the 1H spectra of proteins, and their application to lysozyme. *J. Magn. Reson.* **31**, 459-469 (1978).
15. R. Richarz, K. Wüthrich, NOE difference spectroscopy: A novel method for observing individual multiplets in proton NMR spectra of biological macromolecules. *J. Magn. Reson.* **30**, 147-150 (1978).
16. H. Sierzputowska-Gracz, R. H. Guenther, P. F. Agris, W. Folkman, B. Golankiewicz, Structure and conformation of the hypermodified purine nucleoside wyosine and its isomers: A comparison of coupling constants and distance geometry solutions. *Magn. Reson. Chem.* **29**, 885-892 (1991).
17. J.-P. Desaulniers, H. M. P. Chui, C. S. Chow, Solution conformations of two naturally occurring RNA nucleosides: 3-Methyluridine and 3-methylpseudouridine. *Bioorg. Med. Chem.* **13**, 6777-6781 (2005).
18. Y.-C. Chang, J. Herath, T. H. H. Wang, C. S. Chow, Synthesis and solution conformation studies of 3-substituted uridine and pseudouridine derivatives. *Bioorg. Med. Chem.* **16**, 2676-2686 (2008).
19. C. F. G. C. Geraldes, H. Santos, A. V. Xavier, A proton relaxation study of the conformations of some purine mononucleotides in aqueous solution. *Can. J. Chem.* **60**, 2976-2983 (1982).
20. P. A. Hart, Conformation of mononucleotides and dinucleoside monophosphates. P[H] and H[H] nuclear Overhauser effects. *Biophys. J.* **24**, 833-848 (1978).
21. H. Santos, A. V. Xavier, C. F. G. C. Geraldes, Conformation of purine mononucleotides by H{H} and P{H} nuclear Overhauser effects. *Can. J. Chem.* **61**, 1456-1464 (1983).
22. H. Rosemeyer *et al.*, Syn-anti conformational analysis of regular and modified nucleosides by 1D 1H NOE difference spectroscopy: a simple graphical method based on conformationally rigid molecules. *J. Org. Chem.* **55**, 5784-5790 (1990).
23. H. M. Berman *et al.*, The Protein Data Bank. *Nucleic Acids Res.* **28**, 235-242 (2000).

24. B. R. Miller *et al.*, MMPBSA.py: An efficient program for end-state free energy calculations. *J. Chem. Theory Comput.* **8**, 3314-3321 (2012).
25. S. Kammler, S. Lykke-Andersen, T. H. Jensen, The RNA exosome component hRrp6 is a target for 5-fluorouracil in human cells. *Mol. Cancer Res.* **6**, 990 (2008).
26. L. Arandel *et al.*, Immortalized human myotonic dystrophy muscle cell lines to assess therapeutic compounds. *Dis. Models Mech.* **10**, 487-497 (2017).
27. A. J. Angelbello *et al.*, Precise small-molecule cleavage of an r(CUG) repeat expansion in a myotonic dystrophy mouse model. *Proc. Natl. Acad. Sci. U. S. A.* **116**, 7799-7804 (2019).
28. A. Mankodi *et al.*, Myotonic dystrophy in transgenic mice expressing an expanded CUG repeat. *Science* **289**, 1769-1773 (2000).
29. T. M. Wheeler *et al.*, Reversal of RNA-dominance by displacement of protein sequestered on triplet repeat RNA. *Science* **325**, 336-339 (2009).
30. F. X. Wilson *et al.* Preparation of bibenzo[d]imidazoles and other biheteroaryl compounds as antibacterial agents. WO2010063996A2 (2010).
31. J. Liu, Q. Zhang, Q. Xia, J. Dong, Q. Xu, Synthesis, characterization and properties of polyimides derived from a symmetrical diamine containing bis-benzimidazole rings. *Polym.* **97**, 987-994 (2012).
